# Dark-induced decrease in ascorbate levels in Arabidopsis leaves occurs independently of ascorbate peroxidase and oxidase, recycling enzymes, and senescence signaling

**DOI:** 10.1101/2025.03.30.646231

**Authors:** Tamami Hamada, Kojiro Yamamoto, Akane Hamada, Takanori Maruta

## Abstract

Ascorbate is a key antioxidant that protects plant cells from oxidative damage. While plants actively synthesize ascorbate during the day, its degradation becomes prominent under prolonged dark conditions. Since ascorbate degradation begins with its oxidized form, dehydroascorbate (DHA), this process inherently requires ascorbate oxidation. However, the molecular mechanisms underlying dark-induced ascorbate oxidation and subsequent degradation remain unclear. In this study, we investigated the role of intracellular and extracellular ascorbate redox regulation in controlling this process. Using Arabidopsis knockout mutants for key enzymes involved in ascorbate oxidation and recycling, including ascorbate peroxidase (APX), monodehydroascorbate reductase (MDAR), dehydroascorbate reductase (DHAR), and ascorbate oxidase (AO), as well as NADPH oxidases (rbohD and rbohF), we found that none of these enzymes significantly influenced the dark-induced decrease in ascorbate levels. Notably, ascorbate levels decreased similarly in newly generated multiple mutants, including a quintuple mutant (Δ*dhar pad2 mdar5*), which has severely impaired ascorbate recycling capacity, and the *ao2 rbohD* double mutant, which is strongly expected to exhibit a highly altered apoplastic redox state. Furthermore, we examined the potential involvement of senescence signaling, including ORESARA1 and ethylene signaling components, but found no evidence for their contribution. These findings indicate that the dark-induced decrease in ascorbate levels is not governed by conventional pathways for ascorbate oxidation and recycling or senescence signaling processes, suggesting an unidentified regulatory mechanism.

## 1. Introduction

Leaves contain exceptionally high concentrations of ascorbate, a key antioxidant that plays a crucial role in protecting cells from oxidative damage (Smirnoff, 2018). The regulation of leaf ascorbate levels is strongly influenced by light, with significant increases observed in response to prolonged or high-intensity light exposure. The light-dependent activation of ascorbate biosynthesis primarily drives this process (Smirnoff, 2018; Maruta, 2022). Ascorbate is mainly localized in intracellular compartments, where it scavenges reactive oxygen species (ROS), including the superoxide anion radical, singlet oxygen, hydrogen peroxide, and hydroxyl radical (Smirnoff, 2018). During evolution, plants acquired ascorbate peroxidase (APX), an enzyme that enhances the antioxidant capacity of ascorbate (Maruta *et al*., 2016). Given that ROS production in chloroplasts is inevitable due to photosynthetic activity (Asada, 1999), the ability to accumulate ascorbate and establish ascorbate-dependent antioxidant systems has likely been crucial for mitigating photooxidative stress (Maruta *et al*., 2024).

A rapid decline in leaf ascorbate levels occurs when plants transition from light to dark (Yabuta *et al*., 2007). This reduction is primarily attributed to the inactivation of the major ascorbate biosynthetic pathway, which consists of eight enzymatic steps (Wheeler *et al*., 1998; Smirnoff, 2018). For example, treatment with L-galactono-1,4-lactone (L-GalL), the final precursor of ascorbate biosynthesis, dramatically increases leaf ascorbate levels under light conditions but has little effect in the dark (Yabuta *et al*., 2008). Additionally, multiple biosynthetic enzymes exhibit significant transcriptional or post-translational suppression under dark conditions (Yabuta *et al*., 2007; Wang *et al*., 2013; Aarabi *et al*., 2023; Bournonville *et al*., 2023). This inactivation of biosynthesis appears to trigger ascorbate degradation. Notably, ascorbate is remarkably degraded under prolonged dark conditions (Yabuta *et al*., 2007; Truffault *et al*., 2017), yet the degradation pathway remains incompletely understood (Ford *et al*., 2024). At least two independent pathways have been well characterized in plants. In one pathway, degradation begins with dehydroascorbate (DHA), the oxidized form of ascorbate, undergoing further oxidation, leading to cleavage of the carbon-carbon bond between C_2_ and C_3_ of the ascorbate (C_6_) backbone. This reaction proceeds via oxalyl-threonate as an intermediate and ultimately yields oxalate (C_2_) and L-threonate (C_4_) (Green and Fry, 2005; Dewhirst and Fry, 2018). In the second pathway, DHA undergoes hydrolysis of its lactone ring to form 2,3-diketo-L-gulonate (DKG), which can then be converted into sugar acids (C_5_) or further oxidized in a ROS-dependent manner to generate 2-oxo-L-threo-pentonate (Dewhirst and Fry, 2018; Dewhirst *et al*., 2020). These pathways proceed irreversibly through non-enzymatic reactions, although unidentified enzymes are suspected to accelerate these processes (Green and Fry, 2005). In tomato leaves, dark treatment induces rapid ascorbate degradation within 24 h, accompanied by the accumulation of degradation products such as oxalate and intermediate compounds (Truffault *et al*., 2017).

Since ascorbate degradation originates from DHA, it is expected to be influenced by the ascorbate redox cycling system. Pioneering work using Rosa cell suspension cultures has clearly demonstrated that ascorbate degradation mainly occurs in the apoplast (Green and Fry, 2005). This is consistent with the presence of an unidentified esterase activity in apoplastic fractions that accelerates the oxalyl-threonate pathway (Green and Fry, 2005). If we assume, based on these prior studies, that dark-induced ascorbate degradation primarily occurs in the apoplast, then the involvement of an ascorbate exporter responsible for releasing reduced ascorbate into the extracellular space would be necessary. However, such a transporter has yet to be identified. Moreover, despite the importance of this hypothesis, direct experimental evidence demonstrating that dark-induced ascorbate loss in leaves occurs specifically in the apoplast remains limited. Thus, it remains unclear whether this process takes place intracellularly, extracellularly, or in both compartments.

In the intracellular space, the predominant regulatory system is the ascorbate-glutathione pathway (Asada, 1999; Foyer and Kunert, 2024). In this system, APX reduces hydrogen peroxide to water using reduced ascorbate (ASC) as an electron donor (Maruta *et al*., 2016), thereby oxidizing ASC to monodehydroascorbate (MDHA). Unless rapidly recycled, MDHA undergoes spontaneous disproportionation into DHA and ASC. DHA is highly unstable and can be irreversibly degraded. Monodehydroascorbate reductase (MDAR) and dehydroascorbate reductase (DHAR) function to recycle ASC by reducing MDHA and DHA, using NAD(P)H and reduced glutathione (GSH), respectively (Ding *et al*., 2020; Maruta *et al*., 2024). The DHAR reaction generates oxidized glutathione (GSSG), which is subsequently reduced back to GSH by glutathione reductase (GR) in an NADPH-dependent manner (Foyer and Kunert, 2024). The ascorbate-glutathione pathway is distributed across the cytosol, chloroplasts, and mitochondria, with partial localization in peroxisomes (Maruta *et al*., 2024). From the perspective of ascorbate degradation, APX is expected to promote the process, while ascorbate recycling enzymes (MDAR and DHAR) may exert inhibitory effects.

Despite the importance of these enzymes, their activities under long-term dark conditions remain insufficiently studied. Luschin-Ebengreuth and Zechmann (2016) reported that shading the third rosette leaf of Arabidopsis with aluminum foil led to increased MDAR activity and decreased DHAR activity during the early phase (up to day 2), followed by increased APX activity and decreased GR activity in the later phase (from day 4 onwards). Conversely, Wei *et al*. (2017) found that whole-plant shading for two days did not alter the activities of these enzymes. Interestingly, studies in tomato leaves have shown that neither overexpression nor suppression of MDAR affects dark-induced ascorbate degradation (Truffault *et al*., 2017). However, considering the redundancy of ascorbate recycling systems (Maruta *et al*., 2024), a more detailed analysis is required to clarify the impact of these enzymes on ascorbate degradation. For instance, Arabidopsis possesses three *DHAR* genes, and even a triple mutant (Δ*dhar*) lacking all isoforms exhibits normal ascorbate accumulation, even under stress conditions (Rahantaniaina *et al*., 2017; Terai *et al*., 2020), due to the compensatory non-enzymatic reduction of DHA by GSH. When combined with glutathione-deficient mutations such as *phytoalexin-deficient 2* (*pad2*) or *cadmium-sensitive 2* (*cad2*), the resulting quadruple mutants (Δ*dhar pad2* and Δ*dhar cad2*) completely lose their ability to increase ascorbate levels under high-light stress, leading to the accumulation of degradation products (Terai *et al*., 2020; Hamada *et al*., 2023). Beyond tomato (Truffault *et al*., 2017), the roles of APX and ascorbate recycling enzymes in dark-induced ascorbate decrease remain unexplored.

In the extracellular compartment (apoplast), ascorbate oxidase (AO) is a key enzyme involved in ascorbate redox cycling (Mellidou and Kanellis, 2024). AO catalyzes the one-electron oxidation of ascorbate to MDHA, which, in the absence of MDAR in the apoplast, rapidly disproportionates into DHA and ASC. Since DHAR is also absent in this compartment, DHA may either undergo degradation or be taken up into the cell for reduction (Foyer *et al*., 2020). Additionally, class III peroxidases can utilize ascorbate as an electron donor, contributing to ascorbate oxidation (Kvaratskhelia *et al*., 1997). However, the localization and enzymatic properties of these peroxidases remain largely unclear. Since the lack of AO shifts the apoplastic ascorbate pool toward a more reduced state (Yamamoto *et al*., 2005), AO is considered the primary enzyme driving apoplastic ascorbate oxidation. In leaves, NADPH oxidases, particularly respiratory burst homolog D (rbohD) and rbohF, generate superoxide in the apoplast, which could contribute to ascorbate oxidation under prolonged dark conditions (Miller *et al*., 2009; Suzuki *et al*., 2011).

Ascorbate-deficient mutants have been reported to exhibit accelerated leaf discoloration under prolonged dark conditions (Barth *et al*., 2004; Luschin-Ebengreuth and Zechmann, 2016), suggesting a link between ascorbate degradation and dark-induced senescence. ORESARA1 (ORE1), a plant-specific NAC transcription factor (also known as ANAC092), serves as a master regulator of senescence signaling (Oh *et al*., 1997; Balazadeh *et al*., 2008). Under prolonged dark conditions, PHYTOCHROME B-dependent accumulation of PHYTOCHROME-INTERACTING FACTORS (PIFs) occurs, which induces the transcription of key transcription factors involved in ethylene and abscisic acid signaling, including ETHYLENE INSENSITIVE 3 (EIN3), its closest homolog EIN3-LIKE 1 (EIL1), ABSCISIC ACID INSENSITIVE 5 (ABI5), and ENHANCED EM LEVELS (EEL) (Sakuraba *et al*., 2014; Ueda *et al*., 2020). In the quadruple mutant (*pifQ*) lacking PIF1, PIF3, PIF4, and PIF5, as well as in single or multiple mutants of *ein3*, *eil1*, *abi5*, and *eel*, dark-induced senescence is significantly suppressed (Sakuraba *et al*., 2014; Ueda *et al*., 2020), similar to the *ore1* mutant (Oh *et al*., 1997). The involvement of senescence signaling in regulating ascorbate decrease in the dark remains unclear, but ethylene has been suggested to play a role. Gergoff *et al*. (2010) reported that in ethylene signaling mutants, including *ein3*, ascorbate levels before dark treatment were higher than those in the wild type, and dark-induced ascorbate decrease was also suppressed. However, completely contradictory findings have also been reported. Yu *et al*. (2019) showed that ethylene positively influences ascorbate accumulation, with ethylene signaling mutants exhibiting lower ascorbate levels than the wild type.

In addition to senescence signaling, two regulatory proteins have also been implicated in the control of ascorbate levels. CSN5B, a subunit of the COP9 signalosome (CSN), has been suggested to mediate the decline in ascorbate levels under dark conditions by binding to GDP-D-mannose pyrophosphorylase (GMP), a key enzyme in ascorbate biosynthesis, and promoting its degradation (Wang *et al*., 2013). In the *csn5b* mutant, ascorbate levels are elevated under light and remain stable during dark treatment. However, because dark-induced repression of ascorbate biosynthesis involves multiple steps beyond GMP (Maruta, 2022), it remains uncertain whether loss of CSN5B alone is sufficient to sustain ascorbate biosynthesis in the dark. Another candidate, ascorbic acid mannose pathway regulator 1 (AMR1), is a ubiquitin E3 ligase identified as a negative regulator of ascorbate biosynthesis (Zhang *et al*., 2009). The *amr1* knockout mutant exhibits higher ascorbate levels and transcriptional activation of several biosynthetic genes. Interestingly, AMR1 transcript levels increase during age-related senescence (Zhang *et al*., 2009), suggesting a possible role in suppressing ascorbate biosynthesis during the senescence process, as well as under dark conditions.

Based on previous studies and physiological considerations, we hypothesized that either redox turnover regulation and/or senescence signaling may govern ascorbate decreases in the dark. In this study, we investigated the molecular mechanisms underlying the dark-induced decrease in ascorbate levels in Arabidopsis shoots, using a comprehensive set of mutant lines lacking enzymes involved in ascorbate redox turnover regulation. We further examined the potential involvement of senescence signaling triggered by prolonged darkness. However, none of the genes analyzed showed a detectable contribution, indicating that this process occurs through an as-yet-unknown mechanism.

## 2. Materials and methods

### 2.1. Plant materials and growth conditions

*Arabidopsis thaliana* ecotypes Columbia-0 (Col-0) and Nossen-0 (No-0) were used as wild-type plants. The mutant lines used in this study, listed in **Supplemental Table S1**, were obtained from the Arabidopsis Biological Resource Center or the RIKEN Bioresource Research Center. *sapx tapx* double mutants, as well as Δ*dhar pad2* and *pifQ* quadruple mutants, were generated previously (Sakuraba *et al*., 2014; Terai *et al*., 2020; Kameoka *et al*., 2021). *ao2-1 rbohD* and *ao2-2 rbohD* double mutants were generated by crossing. The *ein2 pif4 pif5*, and *ein3-1 eil1-3* mutants (Ueda *et al*., 2020) were kindly provided by Prof. Makoto Kusaba (Hiroshima University).

Plants were grown on half-strength Murashige and Skoog (MS) medium without sucrose. This was because sucrose serves as a substrate for ascorbate biosynthesis. Thus, the plants were maintained under autotrophic conditions. The plates were incubated in a growth chamber (BiOTRON, LH240S, NK System) under a 16-h photoperiod (100 µmol m^-2^ s^-1^) at 22/20°C and 50% relative humidity. After two weeks, the plates were completely shielded from light by wrapping them at least twice with aluminum foil and incubated in the same growth chamber under dark conditions. Shoots were collected before and after dark treatment for subsequent analyses.

### 2.2. Semi-Quantitative Reverse Transcription Polymerase Chain Reaction (RT-PCR)

Semi-quantitative RT-PCR was performed as described by Terai et al. (Terai *et al*., 2020) using primers listed in **Supplemental Table S2**.

### 2.3. Measurements of Ascorbate, Glutathione, and Pyridine Nucleotides

Ascorbate measurements were performed according to Hamada and Maruta (2024) using an ultra-fast liquid chromatography (UFLC) system equipped with a C18 column. Measurements of glutathione and pyridine nucleotides were conducted as described by Noctor *et al*. (2016) without modifications.

### 2.4. Enzyme Assays

Soluble and membrane-bound APX activities were measured following the method of Kameoka *et al*. (2021). DHAR and MDAR activities were determined using the protocols described by Tanaka *et al*. (2021). GR and CAT activities were measured according to Noctor *et al*. (2016). AO activity was assessed following the method of Ueda *et al*. (2015). For AO extraction, 100 mg of frozen shoot samples were ground in liquid nitrogen and extracted with 1 mL of 0.1 M sodium phosphate buffer (pH 6.5). As in Ueda *et al*. (2015), the buffer ionic strength was sufficient to extract AO from the cell wall, as a subsequent extraction step using the same buffer supplemented with 1 M NaCl retrieved only negligible AO activity in Arabidopsis shoots. The AO assay mixture contained 100 μL of enzyme extract, 800 μL of sodium phosphate buffer (pH 5.6), and 100 μL of 2 mM ASC. The reaction mixture was thoroughly mixed, and the kinetics were recorded at 265 nm (extinction coefficient [ε] = 14.3 mM^-1^ cm^-1^).

### 2.5. **Generation of** Δ*dhar pad2 mdar5* Quintuple Mutants

To generate Δ*dhar pad2 mdar5* quintuple mutants, *mdar5* mutations were introduced into the Δ*dhar pad2* background using the CRISPR/Cas9 system. The pKI1.1R vector (Tsutsui and Higashiyama, 2017), carrying a single-guide RNA (sgRNA) designed to target the *MDAR5* gene, was used for genome editing. The guide sequence (5’-GAAGCGATTCTAGAACCGATCGG-3’) was designed using the CRISPR-P 2.0 design tool (Liu *et al*., 2017). The guide sequence was introduced into the pKI1.1R vector following the protocol of Tsutsui and Higashiyama (2017).

The construct was introduced into Δ*dhar pad2* plants via the floral dip method. T_1_ transformants were selected based on hygromycin resistance. In the T_2_ generation, plants in which the pKI1.1R vector had been excised were identified using red fluorescent protein (RFP) fluorescence as a selection marker (Tsutsui and Higashiyama, 2017). Vector excision was further confirmed by genomic PCR. Genomic DNA from the selected individuals was sequenced to verify CRISPR-induced mutations. As a result, two independent Δ*dhar pad2 mdar5* quintuple mutant lines were successfully established.

### 2.6. Accession Numbers

The accession numbers of genes analyzed in this study are as follows:

**Ascorbate metabolism and redox regulation**: AMR1 (At1g65770), AO1 (At4g39830), AO2 (At5g21100), AO3 (At5g21105), APX1 (At1g07890), APX2 (At3g09640), APX3 (At4g35000), APX5 (At4g35970), DHAR1 (At1g19570), DHAR2 (At1g75270), DHAR3 (At5g16710), MDAR1 (At3g52880), MDAR5 (At1g63940), sAPX (At4g08390), tAPX (At1g77490).

**Senescence and stress response**: CSN5B (At1g71230), EIL1 (At2g27050), EIN2 (At5g03280), EIN3 (At3g20770), ORE1 (At5g39610), PAD2 (At4g23100), PIF1 (At2g20180), PIF3 (At1g09530), PIF4 (At2g43010), PIF5 (At3g59060), RbohD (At5g47910), RbohF (At1g64060).

### 2.7. Data Analyses

Statistical analysis methods are described in the figure legends (Tukey–Kramer test, Dunnett’s test, or Student’s *t*-test). All calculations were performed using at least three independent biological replicates. In all experiments, shoots from at least ten plants were pooled and used as a single biological replicate.

## 3. Results

### 3.1. Dark-induced decrease in ascorbate levels occurs with minimal changes in glutathione, pyridine nucleotides, or regulatory enzymes of the ascorbate redox cycle

To assess the impact of darkness on the ascorbate pool size, two-week-old wild-type *Arabidopsis thaliana* (Col-0) plants were incubated in darkness for seven days. Foliar ascorbate levels declined significantly, reaching approximately 58% and 35% of the initial levels after one and two days of dark treatment, respectively, and continued to decrease progressively thereafter in a time-dependent manner (**Figure 1**). The ascorbate redox state remained high under dark conditions but exhibited a gradual decline with prolonged dark exposure. After seven days of darkness, a statistically significant decrease in the ascorbate redox state was observed compared to the initial value (**Figure 1**).

**Figure 1.**
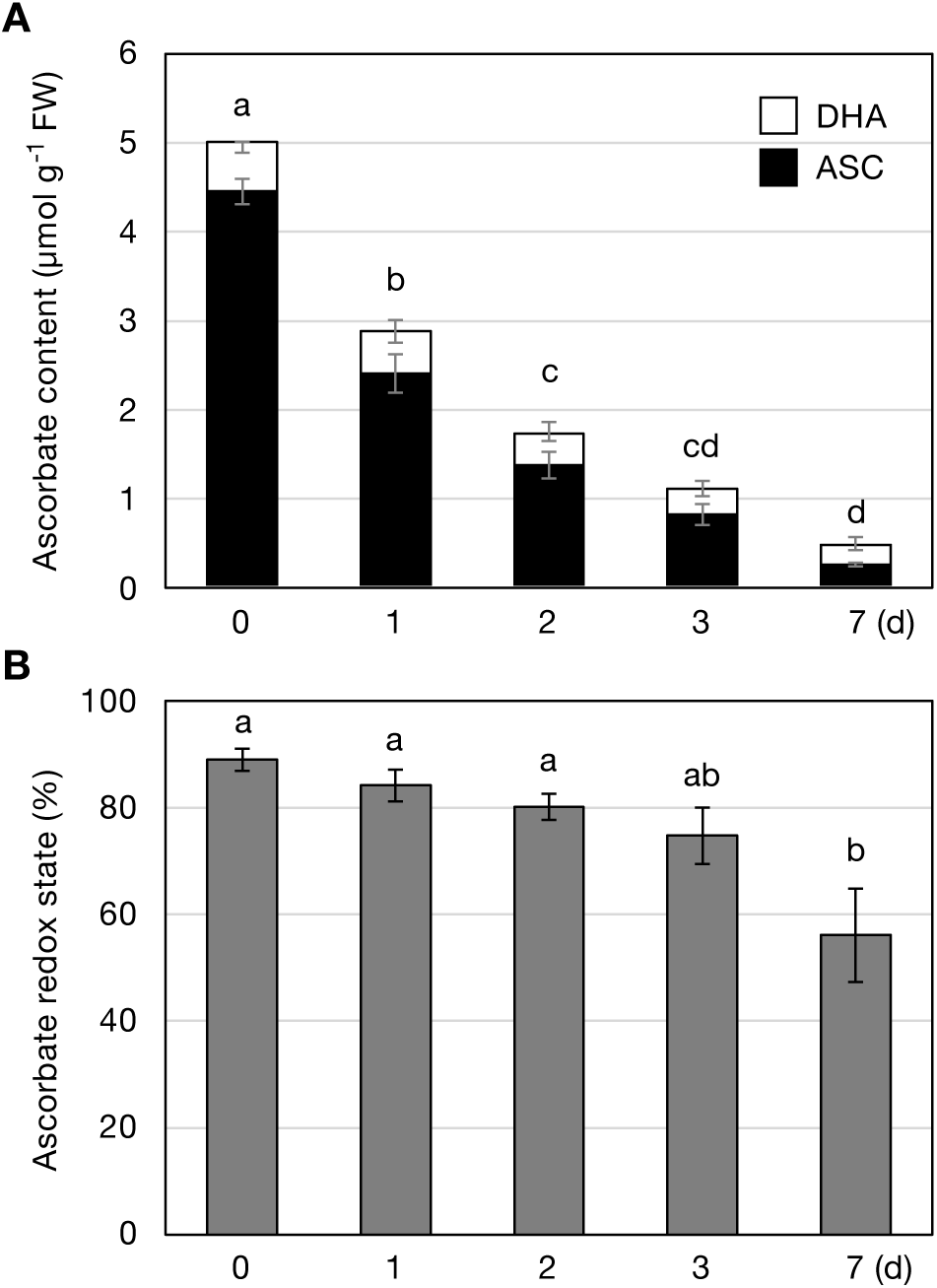
Shoot ascorbate contents in the dark *Arabidopsis thaliana* wild-type plants (Col-0) were grown on half-strength MS medium without sucrose for two weeks and then incubated in darkness for seven days. Shoot ascorbate contents were measured. (A) Total ascorbate content (sum of ASC and DHA). (B) Ascorbate redox state (ratio of ASC to total ascorbate). Data are presented as the mean ± SE of four biological replicates. Different letters indicate significant differences (*P* < 0.05, Tukey–Kramer test). Abbreviations: ASC, reduced ascorbate; DHA, oxidized ascorbate; FW, fresh weight.

Focusing on the first two days, when the most pronounced decrease in ascorbate levels occurred, we measured glutathione and pyridine nucleotide levels, along with the activities of key intracellular enzymes involved in ascorbate redox turnover, including APX, MDAR, DHAR, and GR. The two-day dark treatment had no significant effect on glutathione or NAD(H) levels and their redox states (**Figure 2**). Although a significant decrease in the total NADP(H) pool size was observed under dark conditions, its redox state remained unchanged. Since NADPH indirectly participates in ascorbate recycling by maintaining the glutathione redox state via the GR reaction, the reduction in NADP(H) levels is unlikely to have facilitated ascorbate turnover.

**Figure 2.**
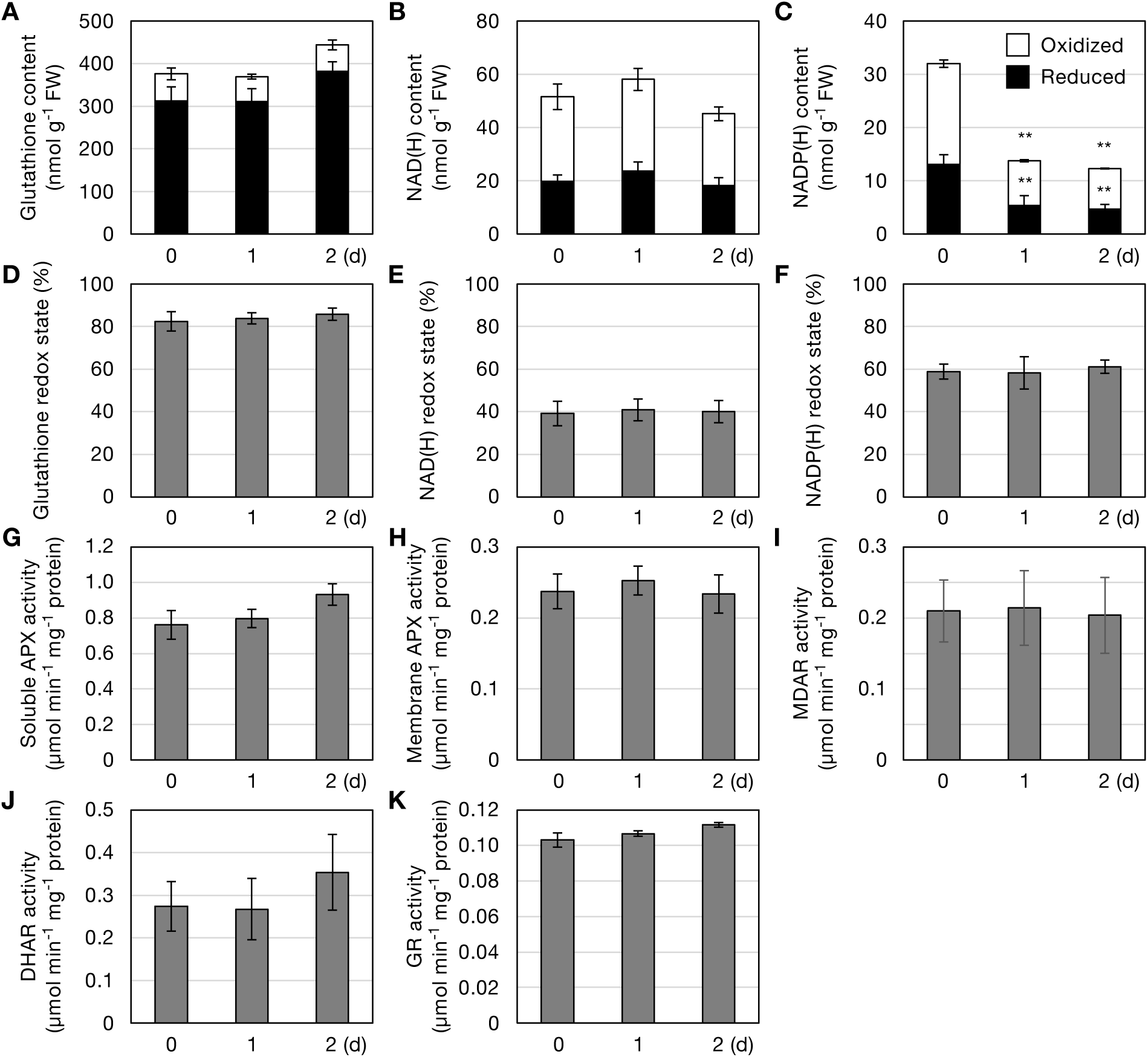
Levels and redox states of glutathione and NAD(P)(H), and activities of enzymes involved in ascorbate redox turnover Two-week-old *Arabidopsis thaliana* wild-type plants (Col-0), grown under the same conditions as in Figure 1, were incubated in darkness for two days. Shoots were collected and analyzed for the following parameters: (A) Total glutathione content (sum of GSH and GSSG). (B) Total NAD(H) content (sum of NADH and NAD_). (C) Total NADP(H) content (sum of NADPH and NADP_). (D) Glutathione redox state (ratio of GSH to total glutathione). (E) NAD(H) redox state (ratio of NADH to total NAD(H)). (F) NADP(H) redox state (ratio of NADPH to total NADP(H)). (G) Soluble APX activity. (H) Membrane-bound APX activity. (I) MDAR activity. (J) DHAR activity. (K) GR activity. Data are presented as the mean ± SE of three or four biological replicates. Significant differences relative to initial values (\*\**P* < 0.01, Dunnett’s test). Abbreviations: FW, fresh weight; GSH, reduced glutathione; GSSG, oxidized glutathione.

APX activity was separately analyzed in water-soluble and insoluble fractions, but no significant changes in activity were detected under dark conditions (**Figure 2**). Similarly, the activities of MDAR, DHAR, and GR remained unchanged during the two-day dark treatment. These findings indicate that Arabidopsis leaf cells maintained a reduced intracellular environment during the initial stages (two days) of dark exposure, with no apparent alterations in the enzymatic activities associated with ascorbate oxidation or recycling.

### 3.2. Impacts of intracellular ascorbate redox cycle regulation on ascorbate decrease in the dark

To examine the role of intracellular ascorbate redox cycling in the dark-induced decrease in ascorbate levels, we analyzed knockout mutants of APX and ascorbate recycling enzymes (MDARs and DHARs). Our initial focus was on APX isoforms, which catalyze ascorbate oxidation. Arabidopsis contains six APX isoforms: two cytosolic (APX1 and APX2), two peroxisomal (APX3 and APX5), and two chloroplastic (sAPX and tAPX). APX1 and APX3 are the predominant isoforms in the cytosol and peroxisomes, respectively (Maruta *et al*., 2016). We used *apx1*, *apx3*, and *apx5* single mutants, as well as *sapx tapx* double mutants, to assess their contributions. APX2 was excluded from this analysis due to its stress-inducible nature (Karpinski *et al*., 1997). All mutants showed reduced APX activity in soluble and/or insoluble fractions compared to wild-type plants (**Supplemental Figure S1**), confirming the roles of these isoforms in intracellular APX activity. When two-week-old wild-type and *apx* mutant plants were incubated in darkness for two days, ascorbate levels declined similarly across all genotypes. The percentage decrease in ascorbate levels relative to pre-treatment levels did not significantly differ between wild-type and mutant plants (**Supplemental Figure S1**).

Next, we examined the ascorbate recycling system. Arabidopsis possesses five MDAR and three DHAR isoforms, alongside a non-enzymatic DHA reduction mechanism involving glutathione. For MDARs, we focused on *mdar1* and *mdar5* mutants, as MDAR1 (localized in the cytosol and peroxisomes) and MDAR5 (localized in chloroplasts and mitochondria) are the primary contributors to measurable MDAR activity in Arabidopsis shoots (Tanaka *et al*., 2021). Consistent with previous reports in tomato plants (Truffault *et al*., 2017), the loss of MDAR did not affect the rate of ascorbate decrease (**Figure 3**). This result was expected, as glutathione-dependent enzymatic (DHAR) and non-enzymatic DHA reduction pathways function complementarily to ensure the robustness of ascorbate recycling (Terai *et al*., 2020; Hamada *et al*., 2023).

**Figure 3.**
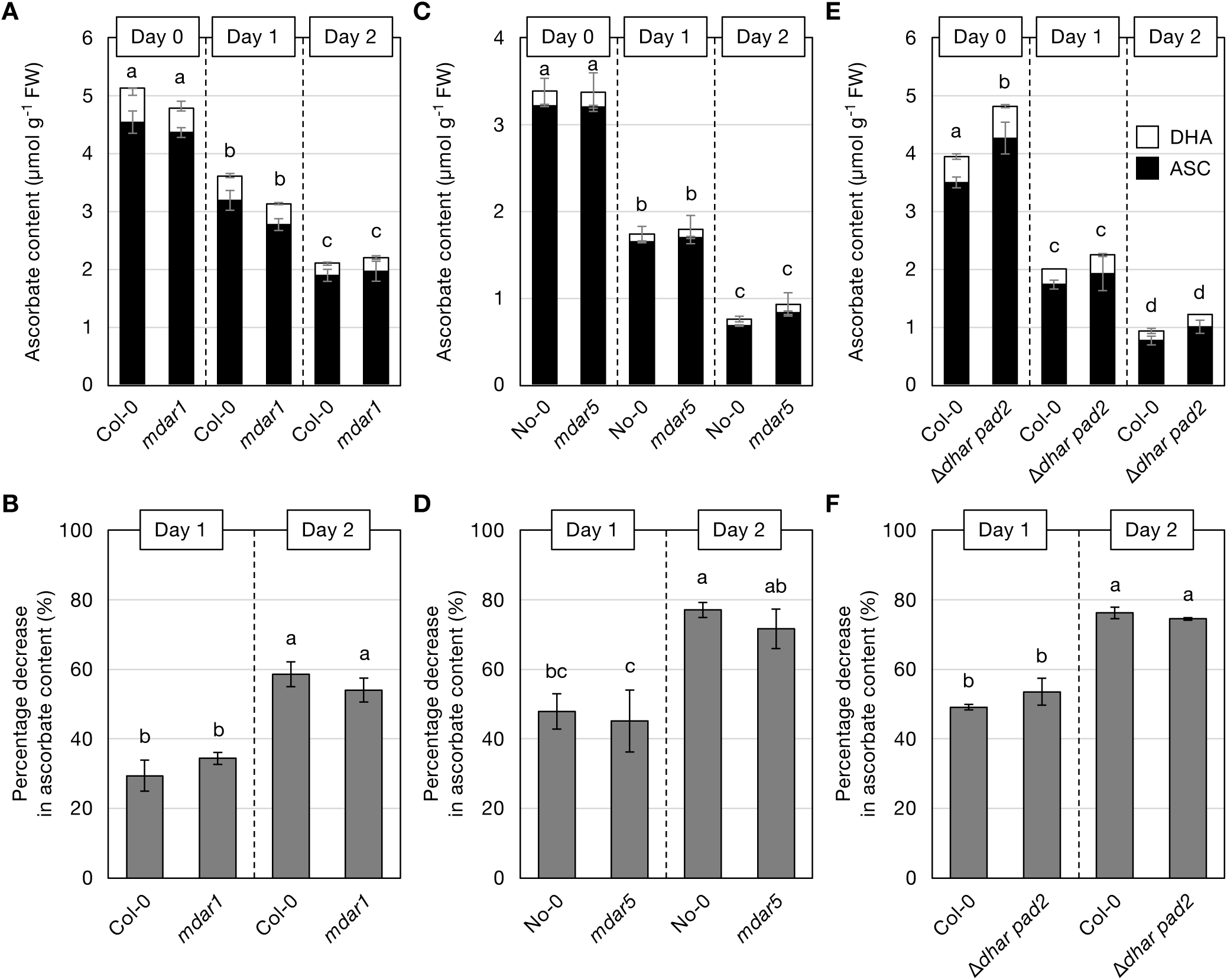
Dark-induced ascorbate decreases in ascorbate recycling mutants Two-week-old *Arabidopsis thaliana* wild-type plants (Col-0 and No-0), *mdar1*, *mdar5*, and Δ*dhar pad2*, grown under the same conditions as in Figure 1, were incubated in darkness for two days. Since the *mdar5* mutant is in the No-0 background, the No-0 ecotype was used for comparison in (C, D), while Col-0 was used as the wild type in all other experiments. (A, C, E) Total ascorbate content (sum of ASC and DHA). (B, D, F) Percentage decrease in ascorbate levels after dark treatment. Data are presented as the mean ± SE of three or four biological replicates. Different letters indicate significant differences (*P* < 0.05, Tukey–Kramer test). Abbreviations: ASC, reduced ascorbate; DHA, oxidized ascorbate; FW, fresh weight.

Therefore, we examined a quadruple mutant (Δ*dhar pad2*) lacking all three *DHAR* genes and deficient in glutathione (via the *pad2* mutation). This mutant exhibits very low ascorbate recycling capacity and is unable to accumulate high ascorbate levels under high-light conditions (Terai *et al*., 2020; Hamada *et al*., 2023). Unlike plants grown in soil (Terai *et al*., 2020; Hamada *et al*., 2023), Δ*dhar pad2* plants grown on MS medium displayed dwarfism and, unexpectedly, higher ascorbate levels than wild-type plants (**Figure 3 and Supplemental Figure S3**). The reason for this increase remains unclear, but growth inhibition may have triggered a stress response, such as the activation of ascorbate biosynthesis. Nevertheless, after dark incubation, ascorbate decrease proceeded similarly in both wild-type and Δ*dhar pad2* plants (**Figure 3**), suggesting that limited ascorbate recycling capacity does not influence ascorbate decrease in the dark.

To further investigate the contribution of ascorbate recycling, we generated additional multiple mutants using the CRISPR-Cas9 system to disrupt the *MDAR5* gene in the Δ*dhar pad2* background, thereby generating quintuple mutants (Δ*dhar pad2 mdar5*). The *MDAR5* gene was targeted using a guide RNA designed to introduce mutations in the second exon (**Figure 4A**), leading to loss of function. Two independent mutant lines were obtained, both harboring frameshift mutations that resulted in premature stop codons. We also attempted to disrupt *MDAR1*, another major isoform in Arabidopsis leaves (Tanaka *et al*., 2021), but have not yet succeeded in generating a knockout mutant. The enzymatic analysis confirmed that MDAR activity in Δ*dhar pad2 mdar* lines was reduced to approximately 64% of the wild-type level, a reduction consistent with that observed in *mdar5* single mutants (**Figure 4B**). DHAR activity remained negligible, similar to the parental Δ*dhar pad2* line (**Figure 4C**). Phenotypically, the quintuple mutants exhibited dwarfism (**Figure 4D and Supplemental Figure S3**), similar to Δ*dhar pad2*, and accumulated higher ascorbate levels than wild-type plants before dark treatment. Ascorbate decrease occurred at a similar rate in both wild-type and Δ*dhar pad2 mdar5* plants (**Figure 4E**). By the second day, a slight trend toward a higher ascorbate depletion rate was observed in the quadruple and quintuple mutants (**Figure 4F**). However, considering the initially higher ascorbate levels in these mutants before dark treatment, this difference was likely due to their elevated ascorbate pool size rather than an intrinsic acceleration of degradation. Notably, focusing on the first day of dark incubation, there was no difference in the percentage decrease in ascorbate levels between the wild-type and mutant plants. We also found that the loss of GR1 did not affect ascorbate decrease (**Supplemental Figure S4**). Overall, these results strongly suggest that the APX and ascorbate recycling systems do not play significant roles in controlling the dark-induced decrease in foliar ascorbate levels.

**Figure 4.**
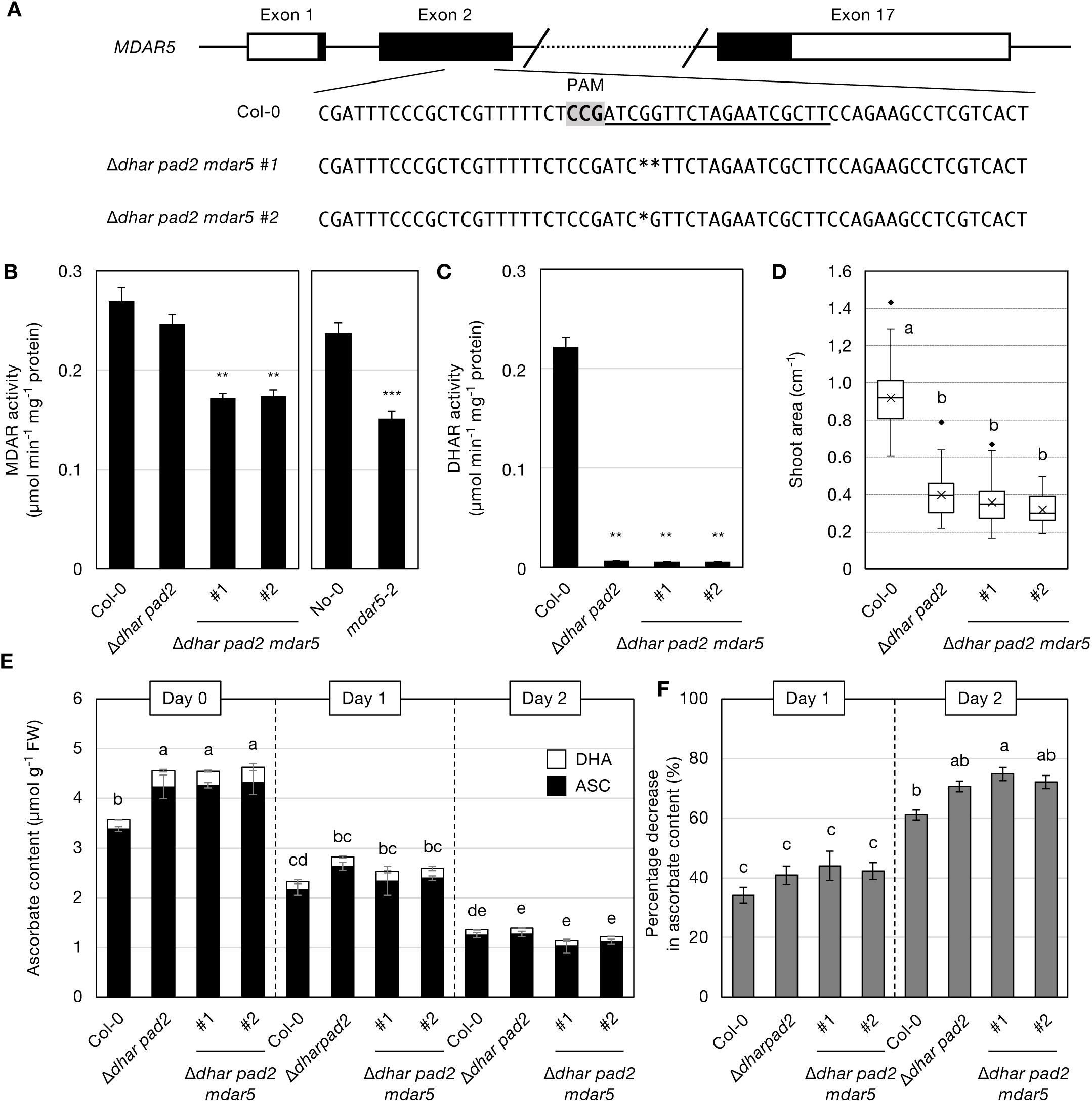
Dark-induced ascorbate decreases in Δ*dhar pad2 mdar5* quintuple mutants (A) Schematic representation of the *MDAR5* gene structure. Black and white boxes indicate exons and untranslated regions (UTRs), respectively, while lines connecting exons indicate introns. The lower panel shows a portion of exon 2, including the guide RNA sequence and the mutations introduced in the genome-edited lines (#1 and #2). (B–F) Two-week-old *Arabidopsis thaliana* wild-type plants (Col-0 and No-0), Δ*dhar pad2*, Δ*dhar pad2 mdar5*, and *mdar5*, grown under the same conditions as in Figure 1, were incubated in darkness for two days. Since the *mdar5* mutant is in the No-0 background, the No-0 ecotype was used for comparison. (B) MDAR activity and (C) DHAR activity before dark treatment. Data are presented as the mean ± SE of three or four biological replicates. Significant differences (\*\**P* < 0.01, Dunnett’s test; \*\*\**P* < 0.001, Student’s t-test). (D) Box plots of shoot area before dark treatment. Data were collected from 35 plants per genotype across three independent experiments. Median and mean values (cross marks) are shown, with boxes representing the 25th and 75th percentiles. Whiskers and diamonds indicate the range and outliers, respectively. (E) Total ascorbate content. (F) Percentage decrease in ascorbate levels after dark treatment. Different letters indicate significant differences (*P* < 0.05, Tukey–Kramer test). Abbreviations: ASC, reduced ascorbate; DHA, oxidized ascorbate; DHAR, dehydroascorbate reductase; FW, fresh weight; MDAR, monodehydroascorbate reductase; PAM, protospacer adjacent motif.

### 3.3. Dark-induced ascorbate decrease in the absence of ascorbate oxidases and/or NADPH oxidases

All enzymes analyzed thus far function within intracellular compartments, whereas AO is localized in the apoplast and operates independently of intracellular ascorbate recycling systems (Mellidou and Kanellis, 2024). Similar to intracellular enzymes, AO activity did not change under dark conditions. Arabidopsis has three genes encoding AO (*AO1*, *AO2*, and *AO3*). Among these, AO2 is the predominant isoform, as indicated by the *ao2-1* mutant, which possesses only ∼20% of the wild-type AO activity (Yamamoto *et al*., 2005). We obtained two independent *ao2* alleles (*ao2-1* and *ao2-2*) (**Supplemental Figure S5**), which exhibited consistently low AO activity both before and after dark treatment (**Figure 5A**). Despite this substantial loss of AO activity, the dark-induced decrease in ascorbate levels was unaffected in both mutants (**Figure 5B and 5C**). We also obtained *ao3* and *ao1* mutants (**Supplemental Figure S5**). The *ao1* knockout did not affect AO activity, whereas *ao3* exhibited a slight (∼21%) but statistically insignificant decrease (**Supplemental Figure S6**). Consistently, neither of these mutants exhibited significant changes in dark-induced ascorbate decrease (**Supplemental Figure S6**).

**Figure 5.**
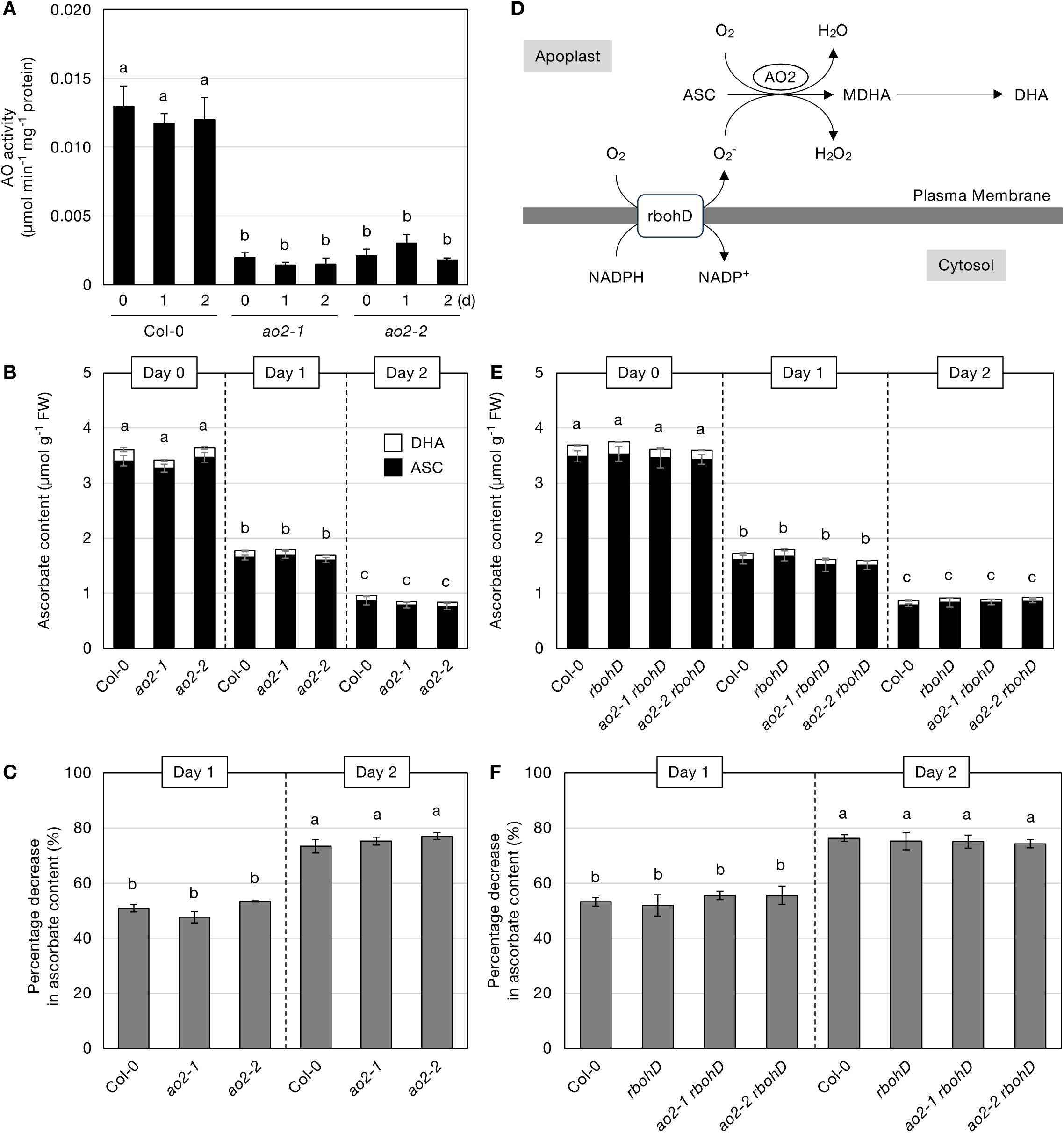
Dark-induced ascorbate decreases in *ao2*, *rbohD*, and *ao2 rbohD* mutants Two-week-old *Arabidopsis thaliana* wild-type plants (Col-0), *ao2-1*, *ao2-2*, *rbohD*, *ao2-1 rbohD*, and *ao2-2 rbohD*, grown under the same conditions as in Figure 1, were incubated in darkness for two days. Shoots were collected and analyzed for (A) AO activity, (B, E) total ascorbate content, and (C, F) percentage decrease in ascorbate levels after dark treatment. (D) Schematic representation of the potential role of NADPH oxidases (RBOHD) in ascorbate oxidation in the apoplast. Data are presented as the mean ± SE of three or four biological replicates. Different letters indicate significant differences (*P* < 0.05, Tukey–Kramer test). Abbreviations: ASC, reduced ascorbate; DHA, oxidized ascorbate; FW, fresh weight; MDHA, monodehydroascorbate.

Since ascorbate oxidation in the apoplast may also be mediated by ROS (**Figure 5D**), we next examined the role of NADPH oxidases. These enzymes, localized to the plasma membrane, generate superoxide by reducing oxygen in the apoplast using intracellular NADPH as an electron donor. In Arabidopsis leaves, rbohD and rbohF are the primary NADPH oxidase isoforms (Miller *et al*., 2009; Suzuki *et al*., 2011). However, dark-induced ascorbate decrease occurred at similar rates in *rbohD* and *rbohF* mutants and wild-type plants (**Figure 5 and Supplemental Figure S7**).

To investigate the potential functional redundancy between AO and rbohD in the apoplast (**Figure 5D**), we generated *ao2-1 rbohD* and *ao2-2 rbohD* double mutants. Given that ascorbate is the primary reductant and RBOHD is the main oxidant-producing enzyme in the apoplast, these double mutants were expected to exhibit strong oxidative imbalances. However, ascorbate levels in the dark decreased at rates comparable to wild-type plants, suggesting that the redox state of the apoplast does not influence ascorbate degradation (**Figure 5D**).

### 3.4. Dark-induced ascorbate decrease is not under the control of senescence signaling

Next, we investigated the regulation of the dark-induced decrease in ascorbate levels from a signaling perspective. Previous studies have reported that CSN5B is required for ascorbate decrease in the dark (Wang *et al*., 2013). Additionally, AMR1, a negative regulator of ascorbate biosynthesis, exhibits increased expression under senescence conditions, where ascorbate levels decline (Zhang *et al*., 2009), suggesting its possible role in regulating the ascorbate pool size in the dark. To explore this, we first compared dark-induced ascorbate degradation in *csn5b* and *amr1* mutants with that in wild-type plants. However, the ascorbate levels in *csn5b* and *amr1* mutants before dark treatment were comparable to those in the wild type. This observation directly contradicts previous reports stating that the loss of CSN5B and AMR1 leads to increased ascorbate levels under normal growth conditions (Zhang *et al*., 2009; Wang *et al*., 2013). Furthermore, we were also unable to reproduce earlier findings that the dark-induced decline in ascorbate levels is completely blocked by the loss of CSN5B (Wang *et al*., 2013). As shown in **Supplemental Figure S8**, the decrease in ascorbate levels during dark in *csn5b* as well as *amr1* mutants during dark treatment was indistinguishable from that in the wild type.

Then, we focused on senescence signaling and examined the role of ORE1, a master regulator of dark-induced senescence signaling, along with its upstream regulators, PIFs. Consistent with previous reports (Oh *et al*., 1997; Sakuraba *et al*., 2014), both *ore1* and *pifQ* mutants exhibited strong suppression of senescence phenotypes under prolonged dark conditions (**Supplemental Figure S9**). Considering that dark-induced senescence typically occurs after extended dark exposure, we tracked temporal changes in ascorbate levels in *ore1* and *pifQ* mutants over a seven-day dark treatment period. However, no significant differences in the rate of ascorbate decrease were observed between these mutants and the wild type (**Figure 6**).

**Figure 6.**
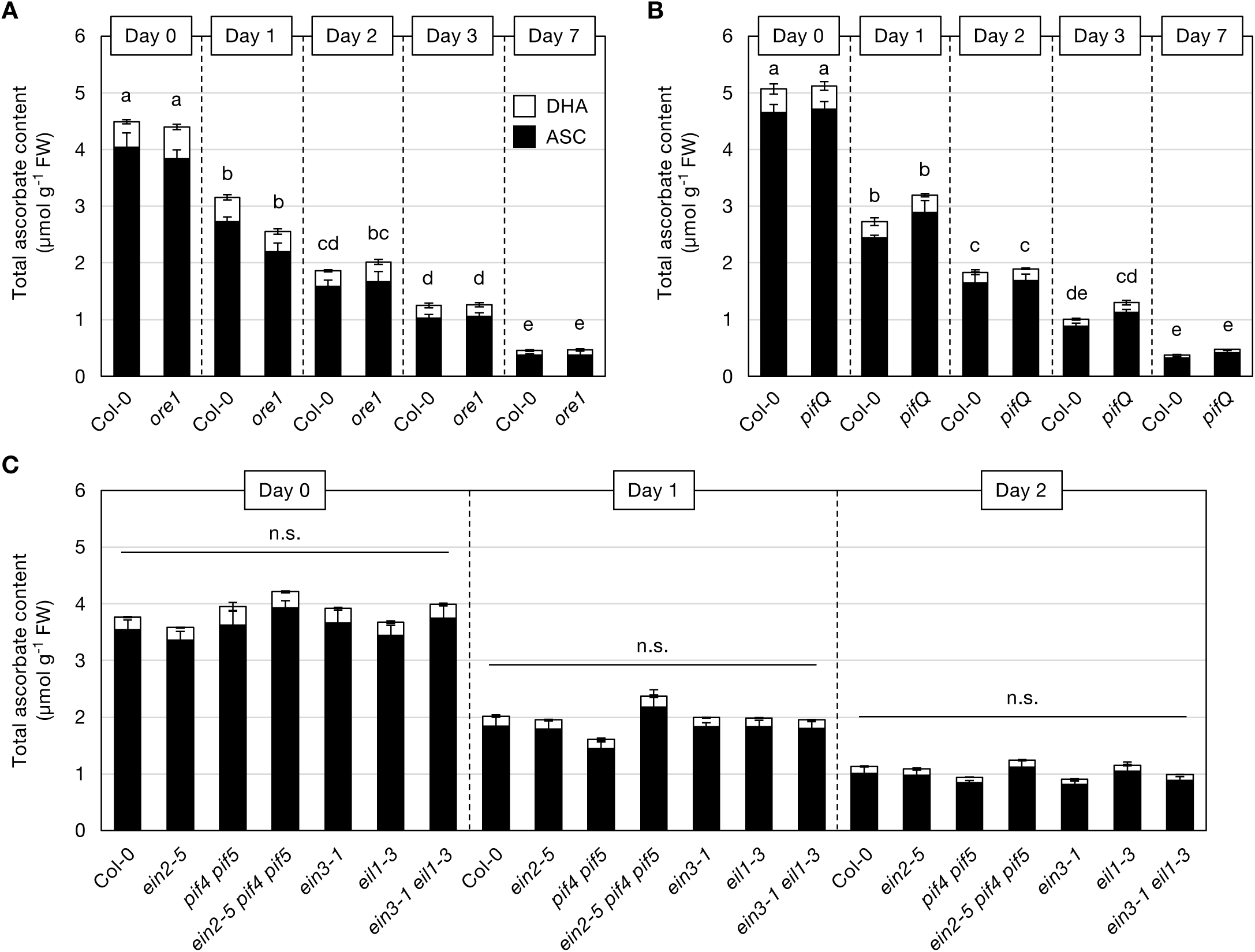
Dark-induced ascorbate decreases in senescence signaling mutants Two-week-old *Arabidopsis thaliana* wild-type plants (Col-0) and senescence-signaling mutants, grown under the same conditions as in Figure 1, were incubated in darkness for two days. Shoots were collected and analyzed for total ascorbate content. (A) *ore1* mutant. (B) *pifQ* mutant. (C) *ein2-5*, *pif4 pif5*, *ein2-5 pif4 pif5*, *ein3-1*, *eil1-3*, and *ein3-1 eil1-3* mutants. Data are presented as the mean ± SE of three or four biological replicates. Different letters indicate significant differences (*P* < 0.05, Tukey–Kramer test). n.s., not significant. The percentage decrease in ascorbate levels for each genotype is shown in Supplemental Figure S10.

We also investigated the potential involvement of ethylene, a key regulator of dark-induced senescence signaling (Liebsch and Keech, 2016). To clarify the relationship between ethylene signaling and dark-induced ascorbate decrease, we analyzed key ethylene signaling mutants, including *ein2*, *ein3*, *eil1*, and the *ein3 eil1* double mutant (Ueda *et al*., 2020). Additionally, we examined the *ein2 pif4 pif5* triple mutant to assess potential interactions between ethylene and PIF-mediated regulatory pathways (Ueda *et al*., 2020). In our study, slight variations in ascorbate levels were observed in some mutants before dark treatment. For example, *ein3 eil1* and *ein2 pif4 pif5* mutants tended to exhibit higher ascorbate levels than the wild type, although these differences were not statistically significant (**Figure 6**). However, upon dark treatment, all mutant lines displayed a decline in ascorbate levels. While *ein3 eil1* and *ein2 pif4 pif5*, which initially had slightly higher ascorbate levels, maintained relatively higher levels after dark treatment, the percentage decrease in ascorbate levels remained comparable to that of the wild type (**Supplemental Figure S10**). Overall, our findings demonstrate that major senescence regulators, including ORE1, PIFs, and ethylene signaling components, do not significantly influence the dark-induced decrease in ascorbate levels. These results suggest that the process operates independently of canonical senescence signaling pathways.

## 4. Discussion

Plants actively synthesize ascorbate during the day to protect cells from photooxidative stress (Maruta *et al*., 2024), while its degradation actively occurs at night. This degradation becomes particularly pronounced under prolonged dark conditions (Truffault *et al*., 2017). Since leaves generally contain high concentrations of ascorbate, they serve as an important dietary source of vitamin C for humans. Thus, dark-induced ascorbate decrease is likely one of the key factors contributing to vitamin C loss during post-harvest storage and distribution. Why and how do plants degrade the ascorbate they accumulate during the day when exposed to shading conditions? Is this merely a passive process resulting from ascorbate turnover? Since ascorbate degradation primarily begins with DHA (Smirnoff, 2018), this process inherently requires ascorbate oxidation. Therefore, it is reasonable to hypothesize that the decrease in ascorbate levels is regulated by the ascorbate redox cycle. In this study, we thoroughly investigated the contribution of intracellular and extracellular redox cycle regulation to the dark-induced decrease in ascorbate levels, as well as its potential association with senescence signaling.

Leaf ascorbate levels declined progressively during dark treatment, with approximately 65% of the initial pool lost within the first two days (**Figure 1**). Despite this rapid decrease, the intracellular redox environment remained highly reduced, as indicated by stable glutathione and pyridine nucleotide levels and redox states (**Figure 2**). Furthermore, no significant changes were detected in the activities of APX, MDAR, DHAR, GR, or AO after two days of dark treatment (**Figure 2**). In contrast, Luschin-Ebengreuth and Zechmann (2016) observed a 50% reduction in DHAR activity and a slight increase in MDAR activity after two days of shading. These discrepancies are likely due to differences in plant age, growth conditions, or experimental setups. However, our findings are consistent with Wei *et al*. (2017), who found that the activities of these enzymes were not altered during whole-plant shading for two days. Thus, under our experimental conditions, two days of dark treatment did not create an oxidative environment in leaf cells.

Despite the absence of an oxidative shift, we hypothesized that the regulation of DHA availability by intracellular or extracellular ascorbate redox cycle enzymes might determine the rate of ascorbate decrease. To test this, we systematically compared dark-induced ascorbate decrease between wild-type plants and mutants deficient in these enzymes. We examined APX, MDAR, and DHAR, which function intracellularly, as well as AO, which operates in the apoplast. Additionally, we analyzed RbohD and RbohF, two major NADPH oxidases involved in ROS production in the apoplast (Miller *et al*., 2009; Suzuki *et al*., 2011). However, using a broad range of single and multiple knockout mutants, we found no evidence that these enzymes contribute to dark-induced ascorbate degradation. These findings strongly suggest that these enzymatic redox systems do not regulate ascorbate decrease under dark conditions.

Our study focused on the major isoforms predominantly expressed in leaves; however, minor isoforms, such as APX2, MDAR2, and MDAR3, that have not yet been analyzed may still contribute to ascorbate metabolism. Notably, recent findings have revealed that cytosolic MDAR2 plays an important role as a major NADPH-dependent MDAR isoform involved in oxidative stress responses (Xu *et al*., 2025). Future investigations may be required to fully exclude their involvement. Nevertheless, data from multiple knockout mutants provide robust support for our conclusions. One particularly informative mutant was the newly generated Δ*dhar pad2 mdar5* quintuple mutant. As discussed earlier, the Δ*dhar pad2* quadruple mutant has severely impaired ascorbate recycling capacity (Terai *et al*., 2020). The additional disruption of the *MDAR5* gene in this background resulted in the Δ*dhar pad2 mdar5* quintuple mutant (**Figure 4**), which is expected to have an even lower ascorbate recycling capacity than the quadruple mutant. However, the dark-induced decrease in ascorbate levels was unaffected in the Δ*dhar pad2 mdar5* quintuple mutant (**Figure 4**).

Another crucial mutant that provided key insights was the *ao2 rbohD* double mutants, which allowed us to assess the role of the extracellular redox environment in ascorbate decrease. Among the three Arabidopsis AO isoforms, AO2 is the predominant one (**Figure 5**). The *ao2* mutant exhibits a markedly reduced apoplastic ascorbate redox state (Yamamoto *et al*., 2005), and AO has been proposed to play a key role in regulating apoplastic redox state in various plant species (Pignocchi *et al*., 2003, 2006; Yamamoto *et al*., 2005; Garchery *et al*., 2013; Karpinska *et al*., 2018). In addition, RbohD and RbohF are the primary NADPH oxidase isoforms responsible for ROS production in Arabidopsis leaves, with RbohD playing a particularly crucial role in apoplastic ROS generation in response to environmental stimuli (Suzuki *et al*., 2011). Previous studies have suggested that prolonged dark exposure may lead to an increase in ROS levels (Wei *et al*., 2017; Tan *et al*., 2020; Hu *et al*., 2021; Jing *et al*., 2022), with NADPH oxidases potentially involved in this process (Jing *et al*., 2022). Based on these findings, the apoplast of *ao2 rbohD* double mutants was expected to be highly reduced. However, the dark-induced decrease in ascorbate levels was completely unaffected in these double mutants. These results indicate that the apoplastic redox state does not regulate the dark-induced decrease in ascorbate levels.

Although our study did not directly investigate the biochemical pathway of ascorbate degradation, it is reasonable to consider that the dark-induced decrease in ascorbate levels is closely associated with ascorbate breakdown. Recently, we identified L-threonate metabolizing domains (*LTD*) as a gene required for the metabolism of L-threonate, a known degradation product of ascorbate (Yamamoto *et al*., 2025). Under the same dark treatment conditions as used in this study, we found that while wild-type plants did not accumulate L-threonate during the decline in ascorbate levels, *ltd* knockout mutants exhibited a marked accumulation of L-threonate (Yamamoto *et al*., 2025). This observation strongly suggests that the dark-induced decrease in ascorbate levels is, at least in part, a result of ascorbate degradation. Thus, our current findings strongly suggest that the dark-induced decrease in ascorbate levels in Arabidopsis leaves occurs independently of major ascorbate redox regulatory systems. This raises a fundamental question: how is ascorbate degraded in the dark? Considering that ascorbate degradation begins with DHA, one possibility is that an unidentified oxidation mechanism independent of the known redox systems initiates this process. Potential candidates include cytochrome *b*_561_ proteins, which mediate redox reactions at the plasma and vacuolar membranes (Gradogna *et al*., 2023), and class III peroxidases, which can exhibit ascorbate peroxidase-like activity in the apoplast or vacuole (Kvaratskhelia *et al*., 1997). If these enzymes are involved, it would be consistent with the pioneering proposal by Green and Fry (2005) that ascorbate degradation predominantly occurs in the apoplast. In such a scenario, an unidentified transporter responsible for exporting ascorbate to the extracellular space would likely play a crucial role. Also, the vacuole, as a major site for the degradation of various biomolecules and organelles, may serve as a primary location for ascorbate breakdown. Recently, a member of the multidrug and toxic compound extrusion (MATE) family, DTX25, has been identified as a vacuolar ascorbate transporter (Hoang *et al*., 2021), making it a promising target for future investigations into ascorbate degradation.

In addition to redox regulation, we investigated the potential role of senescence signaling in controlling ascorbate decrease. Prolonged dark treatment activates senescence pathways through PIFs, ORE1, and ethylene signaling (Oh *et al*., 1997; Sakuraba *et al*., 2014). Notably, a previous study suggested that dark-induced ascorbate degradation is suppressed in ethylene signaling mutants (Gergoff *et al*., 2010). In contrast, another study reported that ethylene positively regulates ascorbate accumulation, as *ein2* mutants exhibit lower ascorbate levels than wild-type plants (Yu *et al*., 2019). Our findings, however, did not support either of these conflicting reports. Instead, we found that ascorbate levels in ethylene signaling mutants remained comparable to those in wild-type plants under both steady-state and dark conditions (**Figure 6**). Ethylene is a key senescence-associated hormone and, thus, its effects on ascorbate metabolism may be influenced by developmental stage or environmental conditions. Therefore, we can conclude that ethylene signaling does not regulate the dark-induced decrease in ascorbate levels, at least in our experimental conditions. Similar to ethylene signaling mutants, *ore1* and *pifq* mutants also showed no differences in ascorbate decrease compared to wild-type plants. Additionally, our findings suggest that the roles of CSN5B and AMR1 in regulating the ascorbate pool size require reconsideration.

In conclusion, our study demonstrates that the dark-induced decrease in ascorbate levels in Arabidopsis occurs independently of major ascorbate redox regulatory systems and senescence signaling pathways. Whether this process is governed by an active or passive mechanism remains unresolved. However, even in a passive mechanism, ascorbate oxidation is likely the initial step that facilitates its loss, as degradation begins with DHA. This strongly suggests the existence of an unidentified regulatory system controlling ascorbate decrease in the dark. Our findings provide significant progress toward understanding the molecular mechanisms underlying dark-induced ascorbate decrease/degradation and offer new directions for future research.

## Data statement

The data underlying this article are available in the article and its online supplementary material.

## Acknowledgments

We would like to thank Prof. Makoto Kusaba for providing the seeds of mutants.

## Author contributions

**Tamami Hamada**: Conceptualization, Methodology, Validation, Investigation, Data Curation, Writing – Review & Editing.

**Kojiro Yamamoto**: Validation, Writing – Review & Editing.

**Akane Hamada**: Methodology, Validation, Investigation, Writing – Review & Editing.

**Takanori Maruta**: Conceptualization, Methodology, Formal analysis, Data Curation, Writing – Original Draft, Visualization, Supervision, Project administration, Funding acquisition.

## Supplemental data

**Figure S1** APX activity in *apx* mutant lines

**Figure S2** Dark-induced ascorbate decreases in *apx* mutants

**Figure S3** Phenotypes of Δ*dhar pad2* and Δ*dhar pad2 mdar5* mutants before dark treatment

**Figure S4** Dark-induced ascorbate decreases in *gr1* mutants

**Figure S5** Expression of genes tested in knockout mutants

**Figure S6** Dark-induced ascorbate decreases in *ao1* and *ao3* mutants

**Figure S7** Dark-induced ascorbate decreases in *rbohF* mutants

**Figure S8** Dark-induced ascorbate decreases in *csn5b* and *amr1* mutants

**Figure S9** Phenotypes of *ore1* and *pifQ* mutants under extended dark conditions

**Figure S10** Percentage decrease in ascorbate levels after dark treatment in senescence mutants

**Table S1** List of primers used

**Table S2** List of mutants used

## References

F. Aarabi, A. Ghigi, M. Wijesingha Ahchige, M. Bulut, P. Geigenberger, H.E. Neuhaus, A. Sampathkumar, S. Alseekh, A.R. Fernie, Genome-Wide Association Study unveils ascorbate regulation by PAS/LOV PROTEIN during high light acclimation, Plant Physiol. (2023). 10.1093/plphys/kiad323.

K. Asada, THE WATER-WATER CYCLE IN CHLOROPLASTS: Scavenging of active oxygens and dissipation of excess photons, Annu. Rev. Plant Physiol. Plant Mol. Biol. 50 (1999) 601–639. 10.1146/annurev.arplant.50.1.601.

S. Balazadeh, D.M. Riaño-Pachón, B. Mueller-Roeber, Transcription factors regulating leaf senescence in *Arabidopsis thaliana*, Plant Biol. (Stuttg.) 10 Suppl 1 (2008) 63–75. 10.1111/j.1438-8677.2008.00088.x.

C. Barth, W. Moeder, D.F. Klessig, P.L. Conklin, The timing of senescence and response to pathogens is altered in the ascorbate-deficient Arabidopsis mutant *vitamin c-1*, Plant Physiol. 134 (2004) 1784–1792. 10.1104/pp.103.032185.

R.A. Dewhirst, S.C. Fry, The oxidation of dehydroascorbic acid and 2,3-diketogulonate by distinct reactive oxygen species, Biochem. J 475 (2018) 3451–3470. 10.1042/BCJ20180688.

R.A. Dewhirst, L. Murray, C.L. Mackay, I.H. Sadler, S.C. Fry, Characterisation of the non-oxidative degradation pathway of dehydroascorbic acid in slightly acidic aqueous solution, Arch. Biochem. Biophys. 681 (2020) 108240. 10.1016/j.abb.2019.108240.

H. Ding, B. Wang, Y. Han, S. Li, The pivotal function of dehydroascorbate reductase in glutathione homeostasis in plants, J. Exp. Bot. 71 (2020) 3405–3416. 10.1093/jxb/eraa107.

C.M. Ford, C. Sweetman, S.C. Fry, Ascorbate degradation: pathways, products, and possibilities, J. Exp. Bot. 75 (2024) 2733–2739. 10.1093/jxb/erae048.

C.H. Foyer, T. Kyndt, R.D. Hancock, Vitamin C in plants: Novel concepts, new perspectives, and outstanding issues, Antioxid. Redox Signal. 32 (2020) 463–485. 10.1089/ars.2019.7819.

C.H. Foyer, K. Kunert, The ascorbate-glutathione cycle coming of age, J. Exp. Bot. 75 (2024) 2682–2699. 10.1093/jxb/erae023.

C. Garchery, N. Gest, P.T. Do, M. Alhagdow, P. Baldet, G. Menard, C. Rothan, C. Massot, H. Gautier, J. Aarrouf, A.R. Fernie, R. Stevens, A diminution in ascorbate oxidase activity affects carbon allocation and improves yield in tomato under water deficit, Plant Cell Environ. 36 (2013) 159–175. 10.1111/j.1365-3040.2012.02564.x.

G. Gergoff, A. Chaves, C.G. Bartoli, Ethylene regulates ascorbic acid content during dark-induced leaf senescence, Plant Sci. 178 (2010) 207–212. 10.1016/j.plantsci.2009.12.003.

A. Gradogna, L. Lagostena, S. Beltrami, E. Tosato, C. Picco, J. Scholz-Starke, F. Sparla, P. Trost, A. Carpaneto, Tonoplast cytochrome b561 is a transmembrane ascorbate-dependent monodehydroascorbate reductase: functional characterization of electron currents in plant vacuoles, New Phytol. 238 (2023) 1957–1971. 10.1111/nph.18823.

M.A. Green, S.C. Fry, Vitamin C degradation in plant cells via enzymatic hydrolysis of 4-O-oxalyl-L-threonate, Nature 433 (2005) 83–87. 10.1038/nature03172.

T. Hamada, T. Maruta, Measurements of ascorbate and dehydroascorbate in plants using high-performance liquid chromatography, Methods Mol. Biol. 2798 (2024) 131–139. 10.1007/978-1-0716-3826-2_8.

A. Hamada, Y. Tanaka, T. Ishikawa, T. Maruta, Chloroplast dehydroascorbate reductase and glutathione cooperatively determine the capacity for ascorbate accumulation under photooxidative stress conditions, Plant J. 114 (2023) 68–82. 10.1111/tpj.16117.

M.T.T. Hoang, D. Almeida, S. Chay, C. Alcon, C. Corratge-Faillie, C. Curie, S. Mari, AtDTX25, a member of the multidrug and toxic compound extrusion family, is a vacuolar ascorbate transporter that controls intracellular iron cycling in Arabidopsis, New Phytol. 231 (2021) 1956–1967. 10.1111/nph.17526.

K. Hu, X. Peng, G. Yao, Z. Zhou, F. Yang, W. Li, Y. Zhao, Y. Li, Z. Han, X. Chen, H. Zhang, Roles of a cysteine desulfhydrase LCD1 in regulating leaf senescence in tomato, Int. J. Mol. Sci. 22 (2021) 13078. 10.3390/ijms222313078.

T. Jing, K. Liu, Y. Wang, X. Ai, H. Bi, Melatonin positively regulates both dark- and age-induced leaf senescence by reducing ROS accumulation and modulating abscisic acid and auxin biosynthesis in cucumber plants, Int. J. Mol. Sci. 23 (2022) 3576. 10.3390/ijms23073576.

T. Kameoka, T. Okayasu, K. Kikuraku, T. Ogawa, Y. Sawa, H. Yamamoto, T. Ishikawa, T. Maruta, Cooperation of chloroplast ascorbate peroxidases and proton gradient regulation 5 is critical for protecting Arabidopsis plants from photo-oxidative stress, Plant J. 107 (2021) 876–892. 10.1111/tpj.15352.

B. Karpinska, K. Zhang, B. Rasool, D. Pastok, J. Morris, S.R. Verrall, P.E. Hedley, R.D. Hancock, C.H. Foyer, The redox state of the apoplast influences the acclimation of photosynthesis and leaf metabolism to changing irradiance, Plant Cell Environ. 41 (2018) 1083–1097. 10.1111/pce.12960.

S. Karpinski, C. Escobar, B. Karpinska, G. Creissen, P.M. Mullineaux, Photosynthetic electron transport regulates the expression of cytosolic ascorbate peroxidase genes in Arabidopsis during excess light stress, Plant Cell 9 (1997) 627–640. 10.1105/tpc.9.4.627.

M. Kvaratskhelia, C. Winkel, R.N. Thorneley, Purification and characterization of a novel class III peroxidase isoenzyme from tea leaves, Plant Physiol. 114 (1997) 1237–1245. 10.1104/pp.114.4.1237.

D. Liebsch, O. Keech, Dark-induced leaf senescence: new insights into a complex light-dependent regulatory pathway, New Phytol. 212 (2016) 563–570. 10.1111/nph.14217.

H. Liu, Y. Ding, Y. Zhou, W. Jin, K. Xie, L.-L. Chen, CRISPR-P 2.0: An improved CRISPR-Cas9 tool for genome editing in plants, Mol. Plant 10 (2017) 530–532. 10.1016/j.molp.2017.01.003.

N. Luschin-Ebengreuth, B. Zechmann, Compartment-specific investigations of antioxidants and hydrogen peroxide in leaves of *Arabidopsis thaliana* during dark-induced senescence, Acta Physiol. Plant 38 (2016) 133. 10.1007/s11738-016-2150-6.

T. Maruta, How does light facilitate vitamin c biosynthesis in leaves?, Biosci. Biotechnol. Biochem. 86 (2022) 1173–1182. 10.1093/bbb/zbac096.

T. Maruta, Y. Sawa, S. Shigeoka, T. Ishikawa, Diversity and evolution of ascorbate peroxidase functions in chloroplasts: More than just a classical antioxidant enzyme?, Plant Cell Physiol. 57 (2016) 1377–1386. 10.1093/pcp/pcv203.

T. Maruta, Y. Tanaka, K. Yamamoto, T. Ishida, A. Hamada, T. Ishikawa, Evolutionary insights into strategy shifts for the safe and effective accumulation of ascorbate in plants, J. Exp. Bot. 75 (2024) 2664–2681. 10.1093/jxb/erae062.

I. Mellidou, A.K. Kanellis, Revisiting the role of ascorbate oxidase in plant systems, J. Exp. Bot. 75 (2024) 2740–2753. 10.1093/jxb/erae058.

G. Miller, K. Schlauch, R. Tam, D. Cortes, M.A. Torres, V. Shulaev, J.L. Dangl, R. Mittler, The plant NADPH oxidase RBOHD mediates rapid systemic signaling in response to diverse stimuli, Sci. Signal. 2 (2009) ra45. 10.1126/scisignal.2000448.

G. Noctor, A. Mhamdi, C.H. Foyer, Oxidative stress and antioxidative systems: recipes for successful data collection and interpretation, Plant Cell Environ. 39 (2016) 1140–1160. 10.1111/pce.12726.

S.A. Oh, J.H. Park, G.I. Lee, K.H. Paek, S.K. Park, H.G. Nam, Identification of three genetic loci controlling leaf senescence in *Arabidopsis thaliana*, Plant J. 12 (1997) 527–535. 10.1046/j.1365-313x.1997.00527.x.

C. Pignocchi, J.M. Fletcher, J.E. Wilkinson, J.D. Barnes, C.H. Foyer, The function of ascorbate oxidase in tobacco, Plant Physiol. 132 (2003) 1631–1641. 10.1104/pp.103.022798.

C. Pignocchi, G. Kiddle, I. Hernández, S.J. Foster, A. Asensi, T. Taybi, J. Barnes, C.H. Foyer, Ascorbate oxidase-dependent changes in the redox state of the apoplast modulate gene transcript accumulation leading to modified hormone signaling and orchestration of defense processes in tobacco, Plant Physiol. 141 (2006) 423–435. 10.1104/pp.106.078469.

M.-S. Rahantaniaina, S. Li, G. Chatel-Innocenti, A. Tuzet, E. Issakidis-Bourguet, A. Mhamdi, G. Noctor, Cytosolic and chloroplastic DHARs cooperate in oxidative stress-driven activation of the salicylic acid pathway, Plant Physiol. 174 (2017) 956–971. 10.1104/pp.17.00317.

Y. Sakuraba, J. Jeong, M.-Y. Kang, J. Kim, N.-C. Paek, G. Choi, Phytochrome-interacting transcription factors PIF4 and PIF5 induce leaf senescence in Arabidopsis, Nat. Commun. 5 (2014) 4636. 10.1038/ncomms5636.

N. Smirnoff, Ascorbic acid metabolism and functions: A comparison of plants and mammals, Free Radical Biology and Medicine 122 (2018) 116–129. 10.1016/j.freeradbiomed.2018.03.033.

N. Suzuki, G. Miller, J. Morales, V. Shulaev, M.A. Torres, R. Mittler, Respiratory burst oxidases: the engines of ROS signaling, Curr. Opin. Plant Biol. 14 (2011) 691–699. 10.1016/j.pbi.2011.07.014.

X.-L. Tan, Y.-T. Zhao, W. Shan, J.-F. Kuang, W.-J. Lu, X.-G. Su, N.-G. Tao, P. Lakshmanan, J.-Y. Chen, Melatonin delays leaf senescence of postharvest Chinese flowering cabbage through ROS homeostasis, Food Res. Int. 138 (2020) 109790. 10.1016/j.foodres.2020.109790.

M. Tanaka, R. Takahashi, A. Hamada, Y. Terai, T. Ogawa, Y. Sawa, T. Ishikawa, T. Maruta, Distribution and functions of monodehydroascorbate reductases in plants: Comprehensive reverse genetic analysis of *Arabidopsis thaliana* enzymes, Antioxidants (Basel) 10 (2021). 10.3390/antiox10111726.

Y. Terai, H. Ueno, T. Ogawa, Y. Sawa, A. Miyagi, M. Kawai-Yamada, T. Ishikawa, T. Maruta, Dehydroascorbate reductases and glutathione set a threshold for high-light-induced ascorbate accumulation, Plant Physiol. 183 (2020) 112–122. 10.1104/pp.19.01556.

V. Truffault, S.C. Fry, R.G. Stevens, H. Gautier, Ascorbate degradation in tomato leads to accumulation of oxalate, threonate and oxalyl threonate, Plant J. 89 (2017) 996–1008. 10.1111/tpj.13439.

H. Tsutsui, T. Higashiyama, PKAMA-ITACHI vectors for highly efficient CRISPR/Cas9-mediated gene knockout in *Arabidopsis thaliana*, Plant Cell Physiol. 58 (2017) 46–56. 10.1093/pcp/pcw191.

H. Ueda, T. Ito, R. Inoue, Y. Masuda, Y. Nagashima, T. Kozuka, M. Kusaba, Genetic interaction among phytochrome, ethylene and abscisic acid signaling during dark-induced senescence in *Arabidopsis thaliana*, Front. Plant Sci. 11 (2020) 564. 10.3389/fpls.2020.00564.

Y. Ueda, S. Siddique, M. Frei, A novel gene, OZONE-RESPONSIVE APOPLASTIC PROTEIN1, enhances cell death in ozone stress in rice, Plant Physiol. 169 (2015) 873–889. 10.1104/pp.15.00956.

J. Wang, Y. Yu, Z. Zhang, R. Quan, H. Zhang, L. Ma, X.W. Deng, R. Huang, Arabidopsis CSN5B interacts with VTC1 and modulates ascorbic acid synthesis, Plant Cell 25 (2013) 625–636. 10.1105/tpc.112.106880.

B. Wei, W. Zhang, J. Chao, T. Zhang, T. Zhao, G. Noctor, Y. Liu, Y. Han, Functional analysis of the role of hydrogen sulfide in the regulation of dark-induced leaf senescence in Arabidopsis, Sci. Rep. 7 (2017) 2615. 10.1038/s41598-017-02872-0.

G.L. Wheeler, M.A. Jones, N. Smirnoff, The biosynthetic pathway of vitamin C in higher plants, Nature 393 (1998) 365–369. 10.1038/30728.

D. Xu, L. Trémulot, Z. Yang, A. Mhamdi, G. Châtel-Innocenti, L. Mathieu, C. Espinasse, F. Van Breusegem, H. Vanacker, E. Issakidis-Bourguet, G. Noctor, Cytosolic monodehydroascorbate reductase 2 promotes oxidative stress signaling in Arabidopsis, Plant Cell Environ. (2025) in press. doi: 10.1111/pce.15488.

Y. Yabuta, T. Maruta, A. Nakamura, T. Mieda, K. Yoshimura, T. Ishikawa, S. Shigeoka, Conversion of L-galactono-1,4-lactone to L-ascorbate is regulated by the photosynthetic electron transport chain in Arabidopsis, Biosci. Biotechnol. Biochem. 72 (2008) 2598–2607. 10.1271/bbb.80284.

Y. Yabuta, T. Mieda, M. Rapolu, A. Nakamura, T. Motoki, T. Maruta, K. Yoshimura, T. Ishikawa, S. Shigeoka, Light regulation of ascorbate biosynthesis is dependent on the photosynthetic electron transport chain but independent of sugars in Arabidopsis, J. Exp. Bot. 58 (2007) 2661–2671. 10.1093/jxb/erm124.

A. Yamamoto, M.N.H. Bhuiyan, R. Waditee, Y. Tanaka, M. Esaka, K. Oba, A.T. Jagendorf, T. Takabe, Suppressed expression of the apoplastic ascorbate oxidase gene increases salt tolerance in tobacco and Arabidopsis plants, J. Exp. Bot. 56 (2005) 1785–1796. 10.1093/jxb/eri167.

K. Yamamoto, Y. Yamashita, T. Hamada, A. Miyagi, H. Murayama, A. Hamada, T. Maruta, Metabolism of L-threonate, an ascorbate degradation product, requires a protein with L-threonate metabolizing domains in Arabidopsis, bioRxiv 2025.05.18.654769 (2025). 10.1101/2025.05.18.654769.

Y. Yu, J. Wang, S. Li, X. Kakan, Y. Zhou, Y. Miao, F. Wang, H. Qin, R. Huang, Ascorbic acid integrates the antagonistic modulation of ethylene and abscisic acid in the accumulation of reactive oxygen species, Plant Physiol. 179 (2019) 1861–1875. 10.1104/pp.18.01250.

W. Zhang, A. Lorence, H.A. Gruszewski, B.I. Chevone, C.L. Nessler, AMR1, an Arabidopsis gene that coordinately and negatively regulates the mannose/l-galactose ascorbic acid biosynthetic pathway, Plant Physiol. 150 (2009) 942–950. 10.1104/pp.109.138453.

