## Supplementary material for "Dark-induced decrease in ascorbate levels in Arabidopsis leaves occurs independently of ascorbate peroxidase and oxidase, recycling enzymes, and senescence signaling": Figures S1-S10

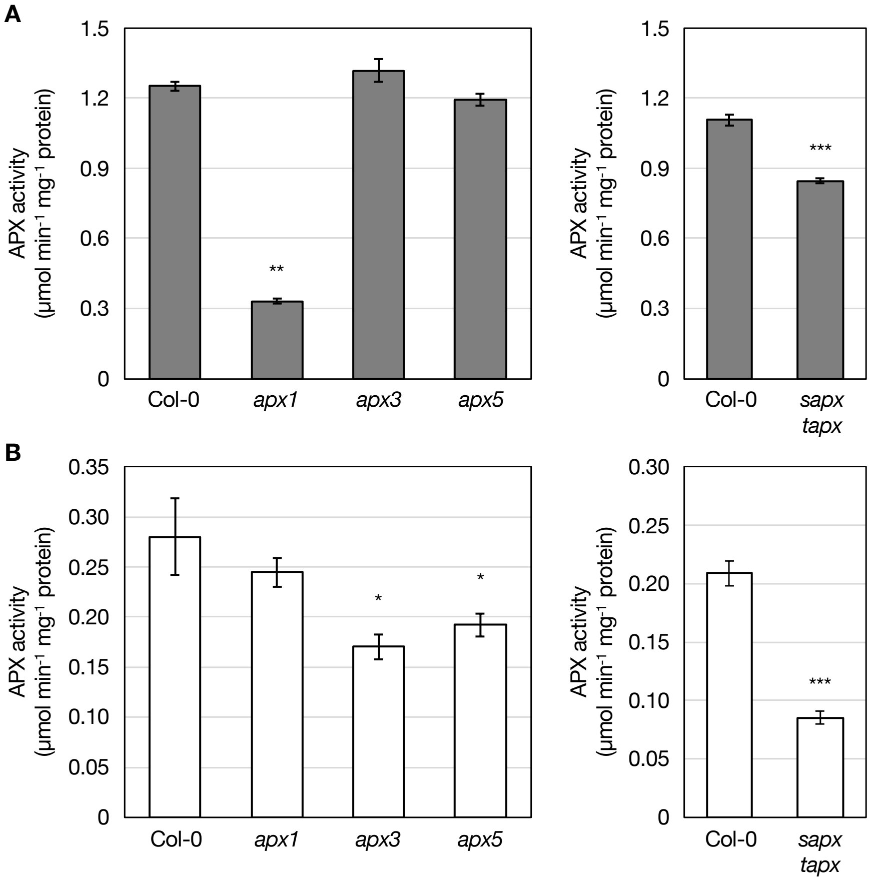


**Supplemental Figure S1** APX activity in *apx* mutant lines

*Arabidopsis thaliana* wild-type plants (Col-0), *apx1*, *apx3*, *apx5*, and *sapx tapx* mutants were grown under the same conditions as in **Figure 1** for two weeks. Shoots were collected and analyzed for APX activity. (A) Soluble APX activity (cytosolic and stromal isoforms). (B) Membrane-bound APX activity (thylakoid and peroxisomal membrane isoforms). Data are presented as the mean ± SE of four biological replicates. Significant differences were determined using Dunnett's test ($^{*}P < 0.05$, $^{**}P < 0.01$) relative to wild-type values. For comparisons between Col-0 and *sapx tapx*, Student's *t*-test was used ($^{***}P < 0.001$). Abbreviation: APX, ascorbate peroxidase.

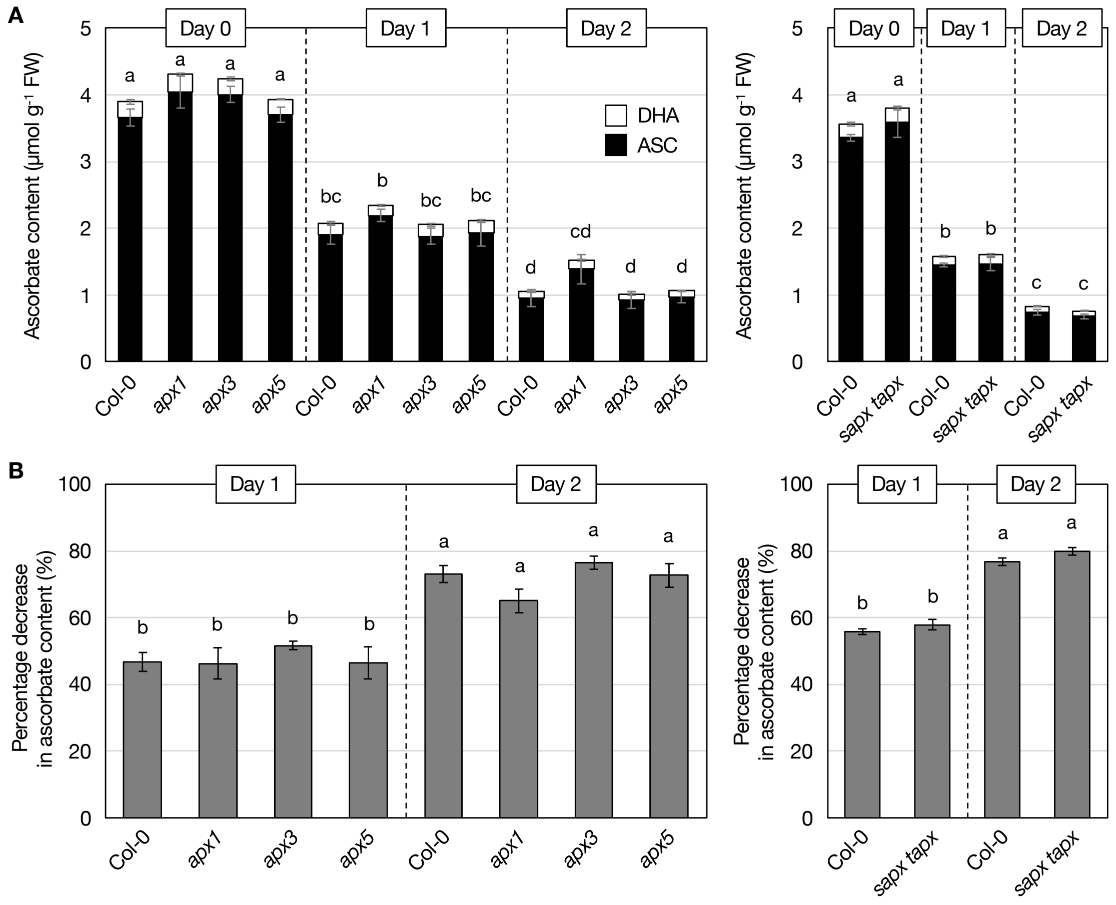


**Supplemental Figure S2** Dark-induced ascorbate decreases in *apx* mutants

Two-week-old *Arabidopsis thaliana* wild-type plants (Col-0), *apx1*, *apx3*, *apx5*, and *sapx tapx* mutants, grown under the same conditions as in **Figure 1**, were incubated in darkness for two days. Ascorbate contents in shoots were measured. (A, C, E) Total ascorbate content (sum of ASC and DHA). (B, D, F) Percentage decrease in ascorbate levels after dark treatment. Data are presented as the mean ± SE of three or four biological replicates. Different letters indicate significant differences ($P < 0.05$, Tukey–Kramer test). Abbreviations: APX, ascorbate peroxidase; ASC, reduced ascorbate; DHA, oxidized ascorbate; FW, fresh weight.

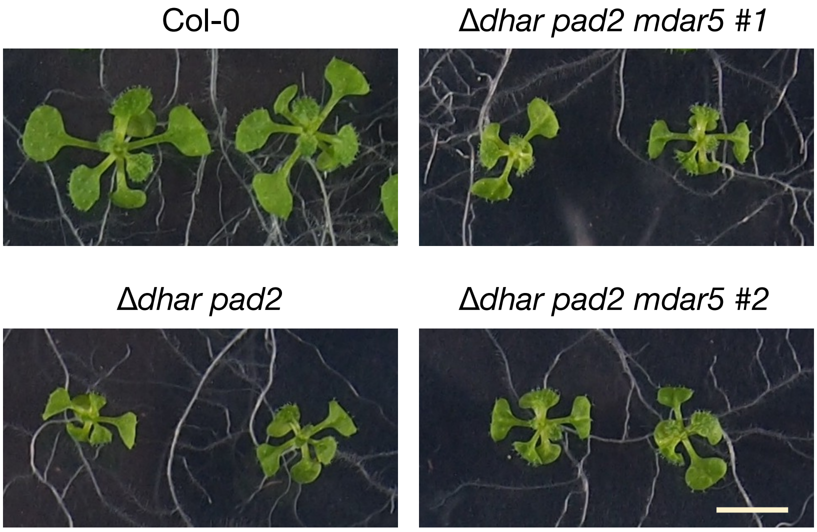


**Supplemental Figure S3** Phenotypes of Δ*dhar pad2* and Δ*dhar pad2 mdar5* mutants before dark treatment

*Arabidopsis thaliana* wild-type plants (Col-0), Δ*dhar pad2*, and Δ*dhar pad2 mdar5* (#1 and #2) were grown under the same conditions as in **Figure 1** for two weeks. Representative images of each genotype are shown. Scale bar = 10 mm.

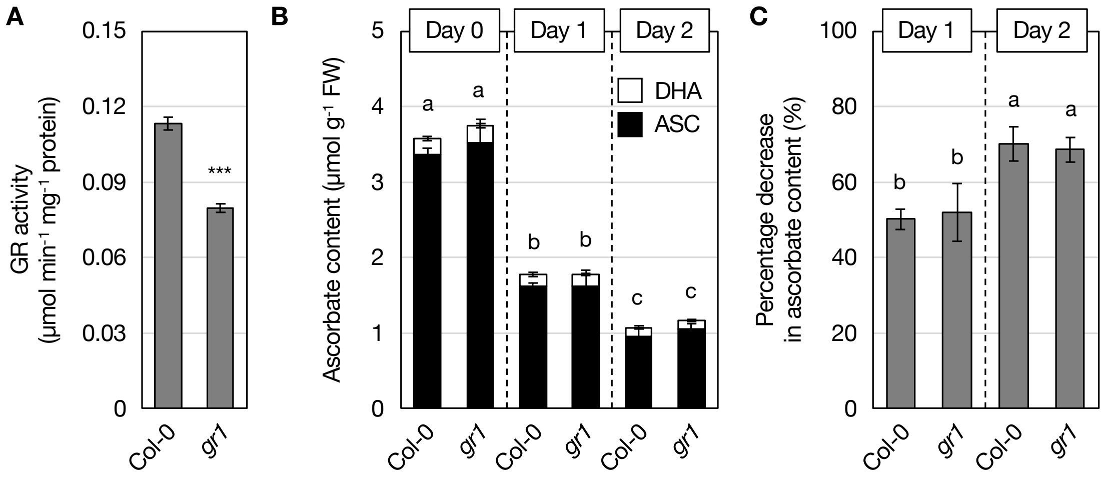


**Supplemental Figure S4** Dark-induced ascorbate decreases in *gr1* mutants

Two-week-old *Arabidopsis thaliana* wild-type plants (Col-0) and *gr1* mutants, grown under the same conditions as in **Figure 1**, were incubated in darkness for two days. Shoots were collected and analyzed. (A) GR activity before dark treatment. (B) Total ascorbate content. (C) Percentage decrease in ascorbate levels after dark treatment. Data are presented as the mean ± SE of four biological replicates. Significant differences were determined using Student's *t*-test ($^{***}P < 0.001$) for (A) and Tukey–Kramer test ($P < 0.05$) for (B, C). Abbreviations: ASC, reduced ascorbate; DHA, oxidized ascorbate; FW, fresh weight; GR, glutathione reductase.

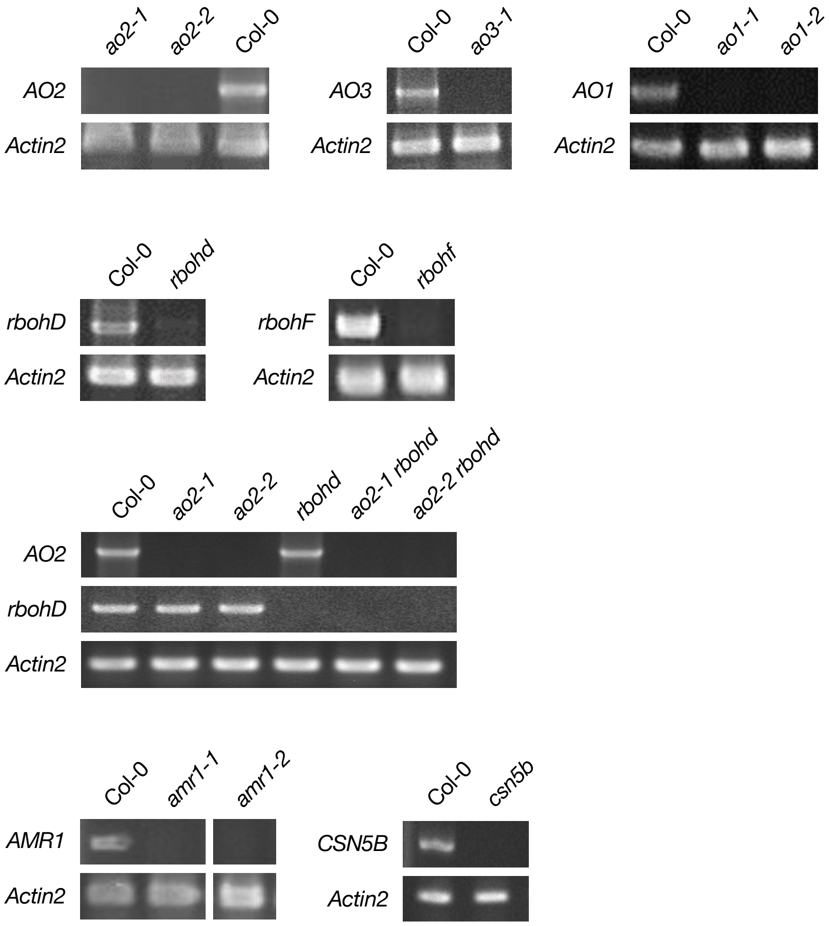


**Supplemental Figure S5** Expression of genes tested in knockout mutants

Shoots from two-week-old plants grown under the same conditions as in **Figure 1** were used for semi-quantitative reverse transcription-PCR.

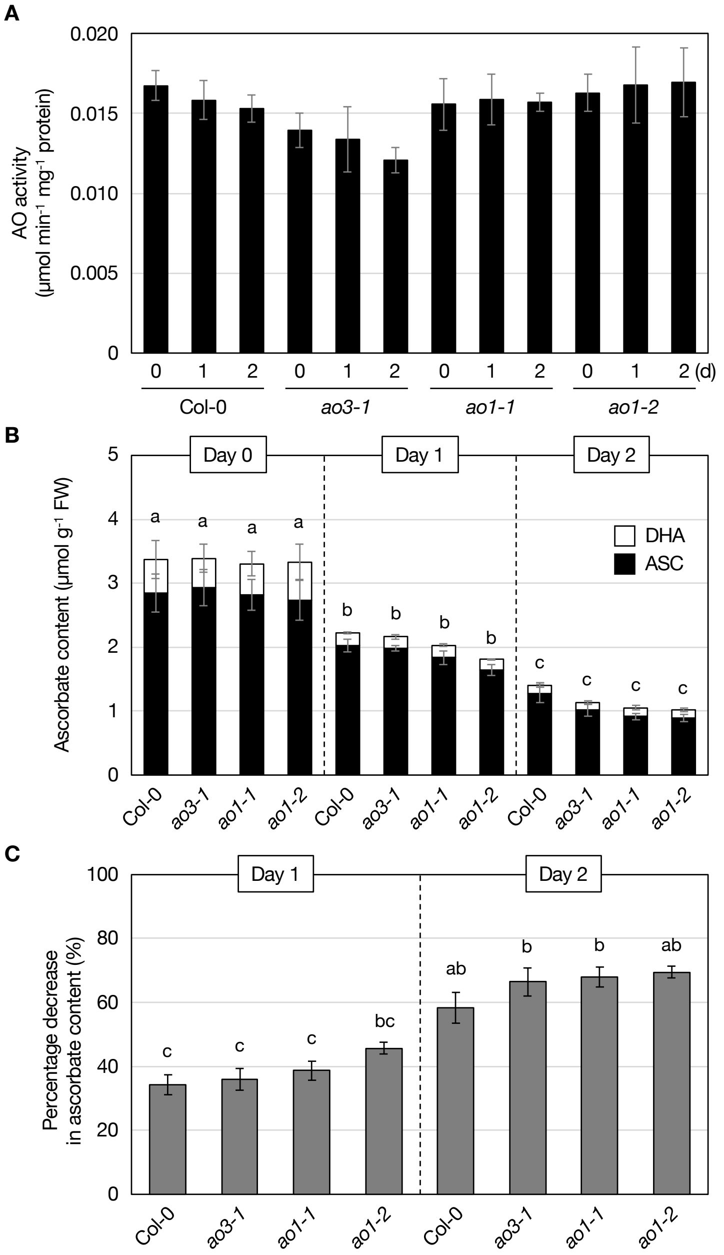


**Supplemental Figure S6** Dark-induced ascorbate decreases in *ao1* and *ao3* mutants

Two-week-old *Arabidopsis thaliana* wild-type plants (Col-0), *ao1-1*, *ao1-2*, and *ao3-1* mutants, grown under the same conditions as in **Figure 1**, were incubated in darkness for two days. Shoots were collected and analyzed. (A) AO activity before dark treatment. (B) Total ascorbate content. (C) Percentage decrease in ascorbate levels after dark treatment. Data are presented as the mean ± SE of three or four biological replicates. Different letters indicate significant differences ($P < 0.05$, Tukey–Kramer test). Abbreviations: AO, ascorbate oxidase; ASC, reduced ascorbate; DHA, oxidized ascorbate; FW, fresh weight.

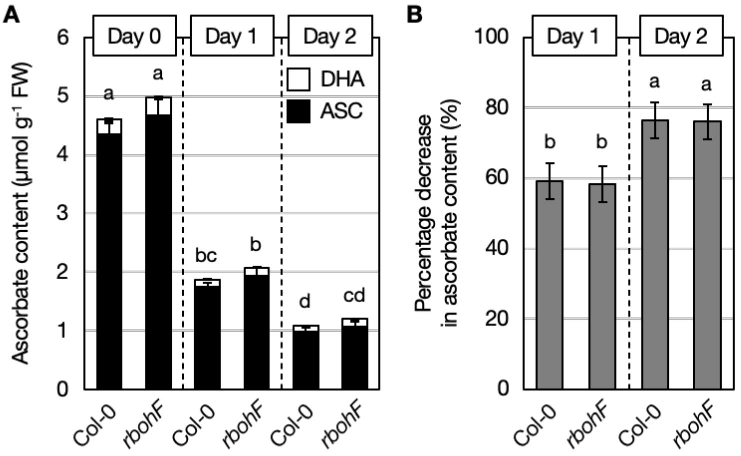


**Supplemental Figure S7** Dark-induced ascorbate decreases in *rbohF* mutants

Two-week-old *Arabidopsis thaliana* wild-type plants (Col-0) and *rbohF* mutants, grown under the same conditions as in **Figure 1**, were incubated in darkness for two days. Ascorbate contents in shoots were measured. (A) Total ascorbate content. (B) Percentage decrease in ascorbate levels after dark treatment. Data are presented as the mean ± SE of three or four biological replicates. Different letters indicate significant differences ($P < 0.05$, Tukey–Kramer test). Abbreviations: ASC, reduced ascorbate; DHA, oxidized ascorbate; FW, fresh weight.

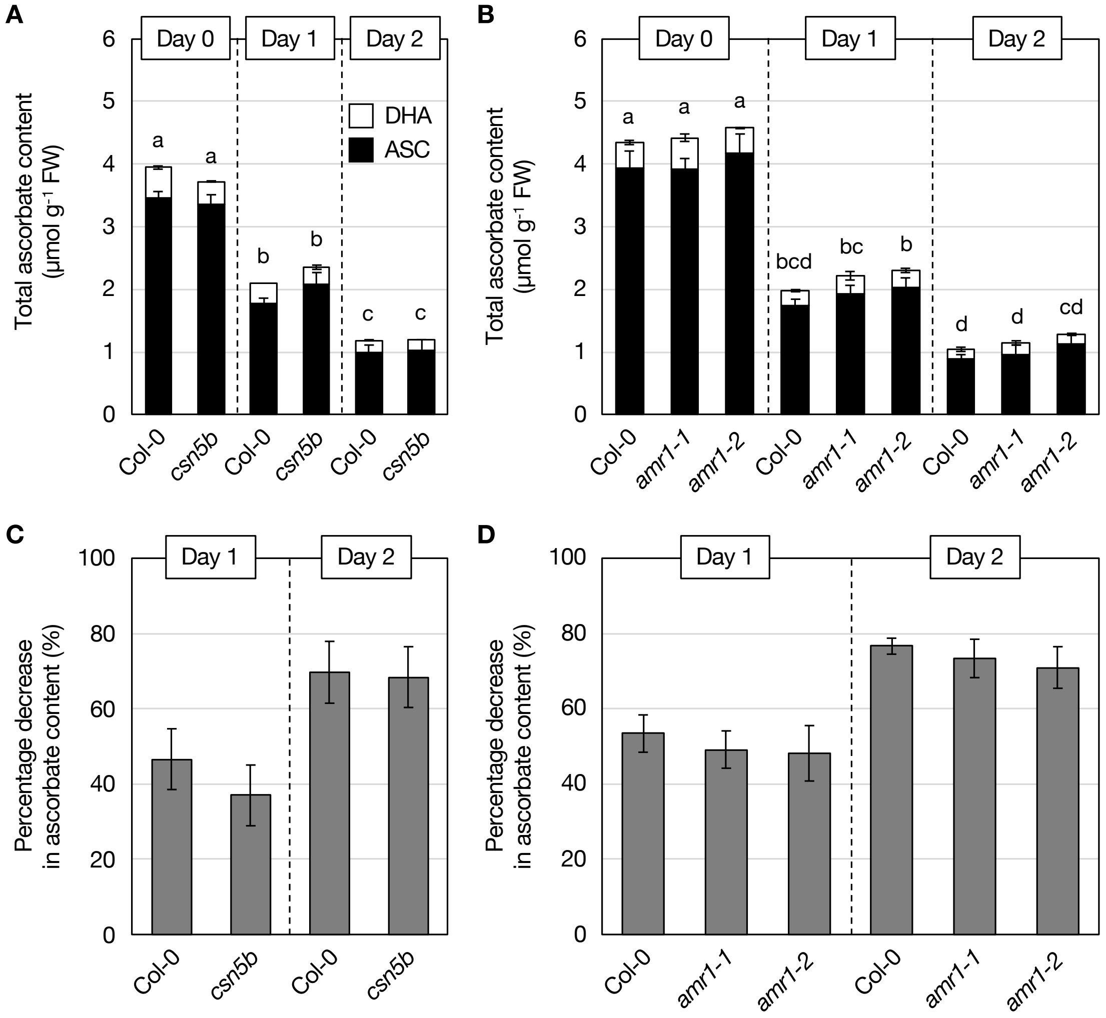


**Supplemental Figure S8** Dark-induced ascorbate decreases in *csn5b* and *amr1* mutants

Two-week-old *Arabidopsis thaliana* wild-type plants (Col-0), *csn5b*, *amr1-1*, and *amr1-2* mutants, grown under the same conditions as in **Figure 1**, were incubated in darkness for two days. Ascorbate contents in shoots were measured. (A) Total ascorbate content. (B) Percentage decrease in ascorbate levels after dark treatment. Data are presented as the mean ± SE of three or four biological replicates. Different letters indicate significant differences ($P < 0.05$, Tukey–Kramer test). Abbreviations: ASC, reduced ascorbate; DHA, oxidized ascorbate; FW, fresh weight.

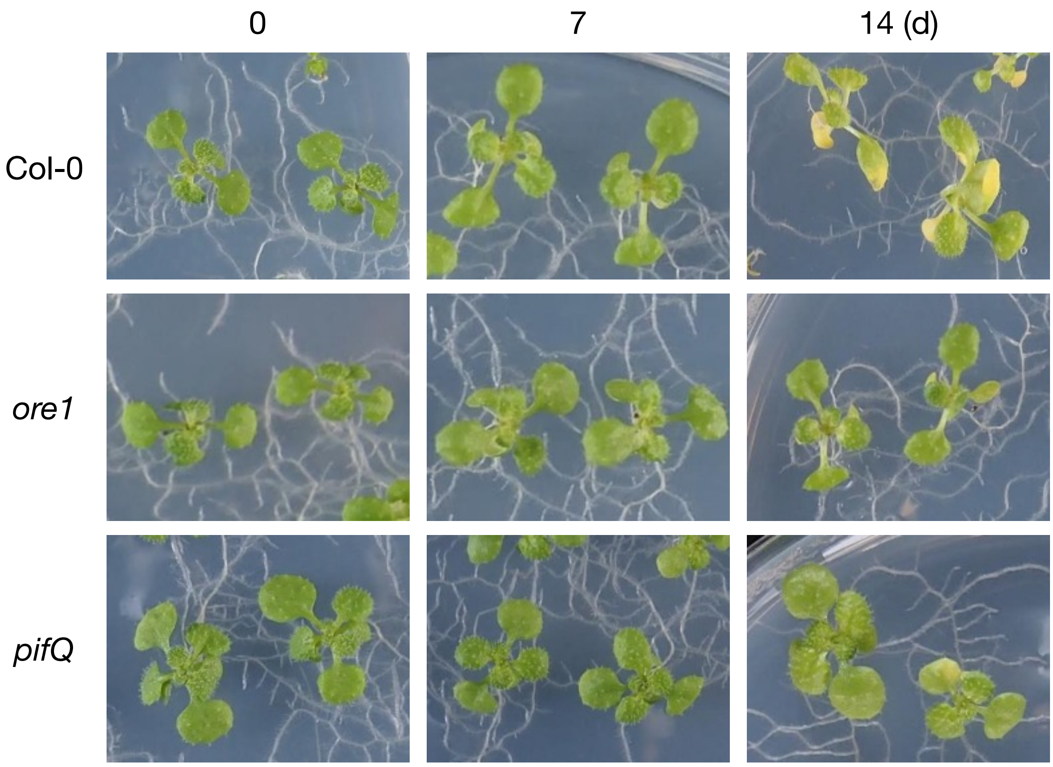


**Supplemental Figure S9** Phenotypes of *ore1* and *pifQ* mutants under extended dark conditions
Two-week-old *Arabidopsis thaliana* wild-type plants (Col-0), *ore1*, and *pifQ* mutants, grown under the same conditions as in **Figure 1**, were incubated in darkness for 14 days. Representative images of each genotype before and after dark treatment are shown.

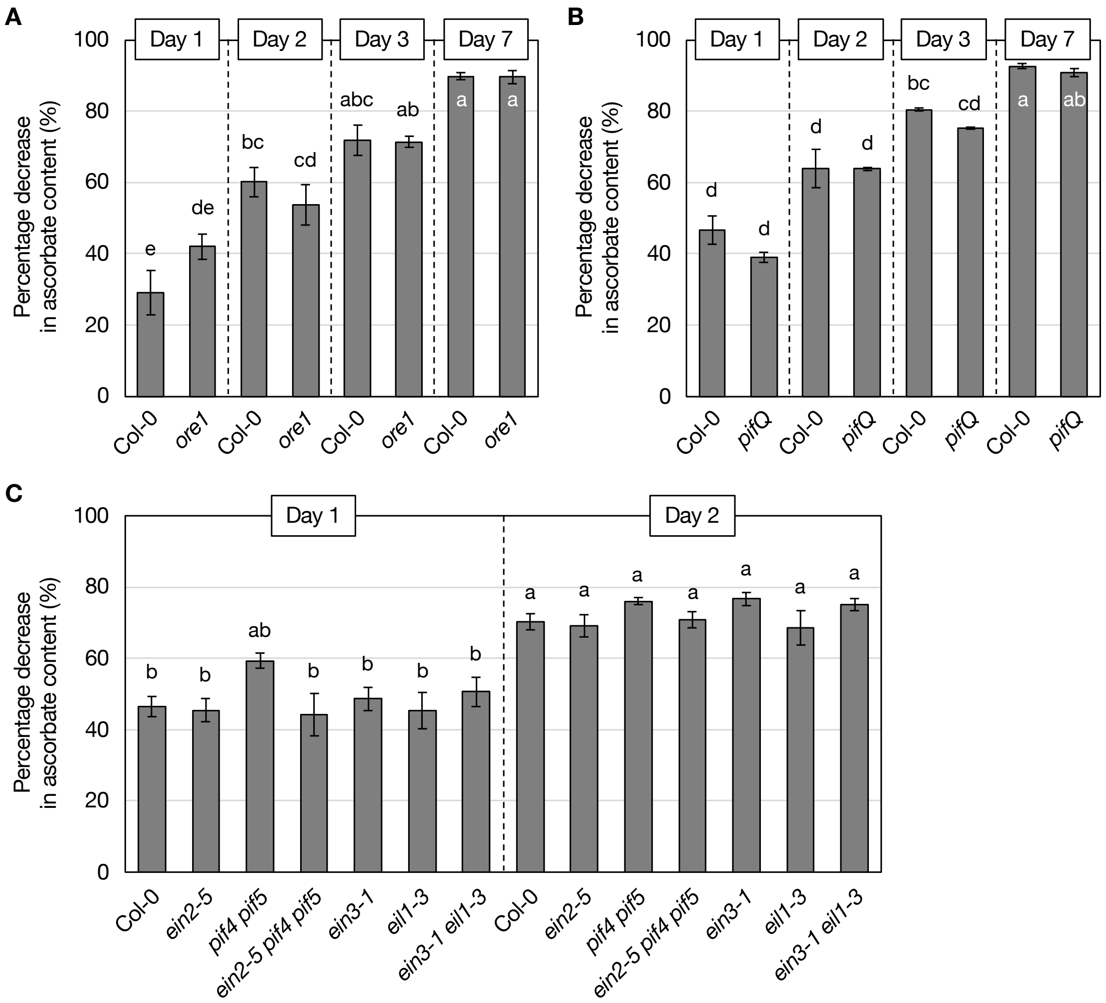


**Supplemental Figure S10** Percentage decrease in ascorbate levels after dark treatment in senescence mutants

The percentage decrease in ascorbate levels after dark treatment in (A) *ore1*, (B) *pifQ*, and (C) *ein2-5*, *pif4 pif5*, *ein2-5 pif4 pif5*, *ein3-1*, *eil1-3*, and *ein3-1 eil1-3* mutants was calculated from the data shown in **Figure 6**. Data are presented as the mean ± SE of three or four biological replicates. Different letters indicate significant differences ($P < 0.05$, Tukey–Kramer test).
