## Supplementary material for "Dark-induced decrease in ascorbate levels in Arabidopsis leaves occurs independently of ascorbate peroxidase and oxidase, recycling enzymes, and senescence signaling": Table S1

**Supplemental Table S1** List of primers used

| Primer | Secence |
|---|---|
| AO1-F | 5’- CCGAAGAGATCATCAGACACAG -3’ |
| AO1-R | 5’- GAGTGGTTAGGTTGTTTGAGC -3’ |
| AO2-F | 5’- ATGGCGGTAATTGTGTGGTG -3’ |
| AO2-R | 5’- CGGTTCATTTAAGGGCGTCC -3’ |
| AO3-F | 5’- GGTGGAGTACAAGTATTGGTCG -3’ |
| AO3-R | 5’- CCAATACGGTTTAATCCCTCC -3’ |
| RbohD-LP | 5’- GTCGCCAAAGGAGGCGCCGA -3’ |
| RbohD-RP | 5’- GGATACTGATCATAGGCGTGGCTCCA -3’ |
| RbohF-F | 5’- GCATCAACTTCACCGGGAA -3’ |
| RbohF-R | 5’- CTATGTAGCCAAGTCTTTCAGG -3’ |
| AMR1-F | 5’- GGCTGATTGGTCTACCTTAC -3’ |
| AMR1-R | 5’- CATGGTATCGTCGGTGAAG -3’ |
| CSN5B-F | 5’- AATGGAGGGTTCGTCGTC -3’ |
| CSN5B-R | 5’- GTCTGGATCCGACGAGT -3’ |
| Actin2 F | 5’- GGCAAGTCATCACGATTGG -3’ |
| Actin2 R | 5’- TCATACTCGGCCTTGGAGATC -3’ |
