## Supplementary material for "Dark-induced decrease in ascorbate levels in Arabidopsis leaves occurs independently of ascorbate peroxidase and oxidase, recycling enzymes, and senescence signaling": Table S2

**Supplemental Table S2** List of mutants used

| Mutant | Allele name/ ABRC or RIKEN BRC stock number |
|---|---|
| *apx1* | SALK_088596 |
| *apx3* | SALK_017480 |
| *apx5* | SALK_144970 |
| *sapx tapx* | *sapx* , SALK_083737; *tapx* , WiscDsLox457-460A17 |
| *mdar1-2* | SALK_034893 |
| *mdar5-2* | 12-4960-1 (RIKEN BRC) |
| *⊿dhar pad2* | *dhar1* , SALK_029966C; *dhar2* , SALK_026089C; *dhar3* , CS820013; *pad2* , CS3804 |
| *gr1* | SALK_105794C |
| *ao1-1* | SALK_136041 |
| *ao1-2* | SALK_098488 |
| *ao2-1* | SALK_108854 |
| *ao2-2* | CS851757 |
| *ao3-1* | SAIL_5_D12 |
| *rbohD* | CS9555 |
| *rbohF* | CS9557 |
| *csn5b* | SALK_036658C |
| *amr1-1* | SALK_113413C |
| *amr1-2* | SALK_081886 |
| *ore1* | SALK_090154 |
| *pifQ* | *pif1-1* , SAIL_256_G07; *pif3-7* , fast neuron mutant; *pif4-2* , SAIL_1288_E07; *pif5-3* , SALK_087012 |
| *ein2-5* | CS16771 |
| *pif4 pif5* | *pif4-102* , SALK_1140393; *pif5-3* , SALK_087012 |
| *ein3-1* | *ein3-1* , CS16710 |
| *eil1-3* | *eil1-3* , SALK_049679 |
